# Herpes simplex virus 1 protein pUL21 stimulates cellular ceramide transport by activating CERT

**DOI:** 10.1101/2022.06.01.494398

**Authors:** Tomasz H. Benedyk, Viv Connor, Eve R. Caroe, Maria Shamin, Dmitri I. Svergun, Janet E. Deane, Cy M. Jeffries, Colin M. Crump, Stephen C. Graham

## Abstract

Herpes simplex virus (HSV)-1 dramatically alters the architecture and protein composition of cellular membranes during infection, but its effects upon membrane lipid composition remain unclear. HSV-1 pUL21 is a virus-encoded protein phosphatase adaptor that promotes dephosphorylation of multiple cellular and virus proteins, including the cellular ceramide transport protein CERT. CERT mediates non-vesicular transport of ceramide from the ER to the *trans*-Golgi network, whereupon ceramide is converted to sphingomyelin and other sphingolipids that play important roles in cell proliferation, cell signalling and membrane trafficking. Using click chemistry to profile the kinetics of sphingolipid metabolism in cultured cells, we show that pUL21-mediated dephosphorylation activates CERT and increases the rate of ceramide to sphingomyelin conversion. Purified pUL21 and full-length CERT interact with sub-micromolar affinity and we map the domains responsible for the interaction. Solving the solution structure of the pUL21 C-terminal domain in complex with the CERT PH and START domains using small-angle X-ray scattering allows us to identify a single amino acid mutation on the surface of pUL21 that disrupts CERT binding *in vitro* and in cultured cells. Sphingolipid profiling demonstrates that ceramide to sphingomyelin conversion is severely diminished in the context of HSV- 1 infection, a defect that is compounded when infecting with a virus encoding the mutated form of pUL21 that lacks the ability to activate CERT. However, virus replication and spread are not significantly altered when pUL21-mediated CERT dephosphorylation is abolished, highlighting that dephosphorylation of other cellular and/or viral targets underpins the important role of pUL21 in HSV-1 biology.

**Significance:** Herpes simplex virus (HSV)-1 causes a life-long dormant infection of neurons, sporadically reactivating to manifest as cold-sores or genital herpes. While the impact of HSV-1 upon the protein content of infected cells has been well studied, we know relatively little about its impact upon cellular lipids. Using bioorthogonal labelling in cultured cells we show that HSV-1 protein pUL21 activates the key cellular lipid transport protein CERT to accelerate the conversion of ceramide to sphingomyelin. HSV-1 infection dramatically alters the kinetics of ceramide metabolism, leading to ceramide accumulation. Mutation of HSV-1 pUL21 to prevent CERT activation further enhances ceramide accumulation but this does not alter the replication or spread of HSV-1, highlighting that other cellular and/or viral proteins represent the critical targets of pUL21-mediated dephosphorylation in cultured cells.

## Introduction

Herpes simplex virus (HSV)-1 is a human pathogen that is estimated to infect the majority of the human population, causing a life-long latent infection [1]. Latent HSV-1 resides in sensory or sympathetic neurons, migrating to the periphery in periodic reactivation events throughout the lifetime of the host. In order to sustain acute (lytic) infection, HSV-1 drastically modifies infected cells [2–5]. In particular, the virus extensively remodels the composition and architecture of cellular membranes to facilitate virus assembly and spread. Nascent capsids leave the nucleus via sequential envelopment and de-envelopment at the inner and outer nuclear membranes [6]. These cytosolic capsids acquire a proteinaceous layer termed ‘tegument’ and bud into the lumen of post-Golgi membranes that are studded with viral glycoproteins (so called ‘secondary envelopment’) [7]. The resultant virus-containing vesicles are transported to cell contact sites where they release the mature virus particles to disseminate the infection [8].

HSV-1 pUL21 is a tegument protein that is conserved in all alphaherpesviruses [9]. This multifunctional protein and its homologues are known to interact with multiple cellular and viral partners, including pUL16 [10, 11], pUL11 [12], gE [12], tubulin [13] and Roadblock-1 [14], and it has been implicated in a number of important processes including capsid nuclear egress [15], viral cell-to-cell spread [12, 16], and retrograde transport along axons to the neuronal cell bodies where latency is established [14, 17]. Mutant HSV-1 lacking pUL21 expression exhibits a 10- to 100-fold replication defect and severely impaired cell-to-cell spread, both of which can be at least partially ascribed to the phosphomodulatory role of pUL21 as a protein phosphatase 1 (PP1) adaptor [18]. PP1 is a highly active and abundant cellular phosphatase [19] and pUL21 recruits PP1 to promote dephosphorylation of multiple substrates, including the viral protein pUL31 that is implicated in viral nuclear egress [18, 20] and the cellular protein CERT that regulates sphingomyelin metabolism [18].

The cytoplasmic ceramide transport protein CERT (a.k.a. Goodpasture’s antigen binding protein, GPBP, encoded by the gene *COL4A3BP*) mediates the non-vesicular trafficking of ceramide from the endoplasmic reticulum (ER) to the *trans*-Golgi network (TGN) and, in doing so, defines the rate of sphingomyelin synthesis [21–23]. CERT contains two well-folded globular domains: an N-terminal pleckstrin homology (PH) domain that mediates its interaction with TGN membranes by binding phosphatidylinositol 4-phosphate, and a C-terminal steroidogenic acute regulatory related lipid transfer (START) domain that directly binds ceramide to mediate its transfer [23]. Crystal structures of both domains have been solved [24, 25]. The ‘middle region’ (MR) that connects the PH and START domains of CERT does not adopt a globular fold, instead containing a coiled-coil region that is likely to mediate CERT self-association [26], a ‘two phenylalanines in an acidic tract’ (FFAT) motif that recruits CERT to the ER membranes via binding proteins VAPA and VAPB [27], and a serine-rich motif (SRM), hyperphosphorylation of which represses CERT activity [28]. In cultured HeLa cells the vast majority of CERT is in an inactive, hyperphosphorylated state (CERT^P^), available to be mobilised via dephosphorylation to increase the rate of ER-to-TGN ceramide transport in response to stimuli such as sphingomyelin depletion [28].

Ceramide (Cer) and sphingolipids like sphingomyelin (SM) are essential for mammalian cell growth and they play important roles in cell signalling, apoptosis and membrane trafficking [29]. Furthermore, sphingolipids and cholesterol participate in the formation of membrane microdomains (including ‘lipid rafts’) that compartmentalise the lipid and protein composition of cellular membranes [30]. This compartmentalisation is especially important in highly polarised cells such as neurons [31]. Although sphingolipids are crucial to host-cell biology, relatively little is known about the interaction between pathogens and cellular sphingolipid metabolism. The bacteria *Chlamydia trachomatis* is known to directly recruit CERT to bacteria-containing intracellular inclusions, thereby increasing the abundance of SM in the inclusion membrane [32], and CERT activity is necessary for the biosynthesis of double-membrane vesicles that serve as the sites of hepatitis C virus and polio virus replication [33]. However, to the best of our knowledge, the binding of CERT to HSV-1 pUL21 [18] is the only known example of a direct interaction between CERT and a virus protein.

Previous studies have shown that HSV-1 infection increases Cer synthesis [34] and that depletion of sphingomyelinase causes a >30-fold decrease in HSV-1 replication [35]. Sphingomyelinase treatment of cultured epithelial cells has only a very modest effect upon HSV-1 entry [36], suggesting that SM is not absolutely required for virus infection. In contrast, recent studies in macrophages demonstrated that acid ceramidase activity is potently antiviral, helping sequester incoming virus particles by promoting their neutralising association with sphingosine-rich intraluminal vesicles within endocytic multi-vesicular bodies [37]. Furthermore, siRNA depletion of CERT has been shown to promote secretion of usually cell-associated HSV-1 virions [38]. It is therefore increasingly clear that cellular sphingolipid metabolism is important for HSV-1 biology, and that further studies are required to obtain a full picture of how specific sphingolipids support and/or prevent HSV-1 infection.

We sought to define the functional consequences of the pUL21:CERT interaction for HSV-1 infection. Using click-chemistry we demonstrated that pUL21-mediated CERT dephosphorylation increases the rate of Cer to SM conversion in cultured cells. We characterised the solution structure of the pUL21 C-terminal domain in complex with the PH and START domains of CERT, identifying a specific pUL21 amino acid required for the interaction. Functional characterisation of mutant HSV-1 encoding this mutated form of pUL21 confirmed that pUL21-mediated CERT dephosphorylation alters the rate of SM synthesis in infected cells. Furthermore, we observed a dramatic increase in the rate of Cer synthesis in infected cells, which was only partly counteracted by pUL21-stimulated CERT activity, but this increase in Cer abundance does not significantly alter virus replication or spread.

## Results

Human cells typically maintain lipidome homeostasis via robust feedback and control mechanisms, which respond to perturbations via local activation of specific signalling pathways [39, 40]. We hypothesised that pUL21, expressed at late points of infection [3], would likely alter the rate of sphingolipid synthesis and thereby affect lipid-mediated signalling events, rather than altering the overall steady-state lipid composition of infected cells. Synthetic lipids with alkyne-containing acyl chains are efficiently processed by mammalian lipid-modifying enzymes and represent powerful tools to probe lipid metabolism [41]. The impact of pUL21-directed CERT dephosphorylation on sphingolipid metabolism was therefore probed using a clickable analogue of sphingosine (Sph), alkyne-Sph, to monitor the rate of sphingomyelin biogenesis in immortalized human keratinocyte (HaCaT) cells. Exogenous Sph is efficiently incorporated into cellular metabolic pathways, being rapidly converted into ceramide (Cer) and then sphingomyelin (SM) or hexosylceramides (HexCer) like glucosylceramide or galactosylceramide [42]. It is also converted into phosphatidylcholine (PC) via a so-called ‘salvage’ pathway (Fig 1A) that directs sphingosine to palmitoyl-CoA which serves as substrate for re-acetylation of lysophosphatidylcholine [43].

**Figure 1.**
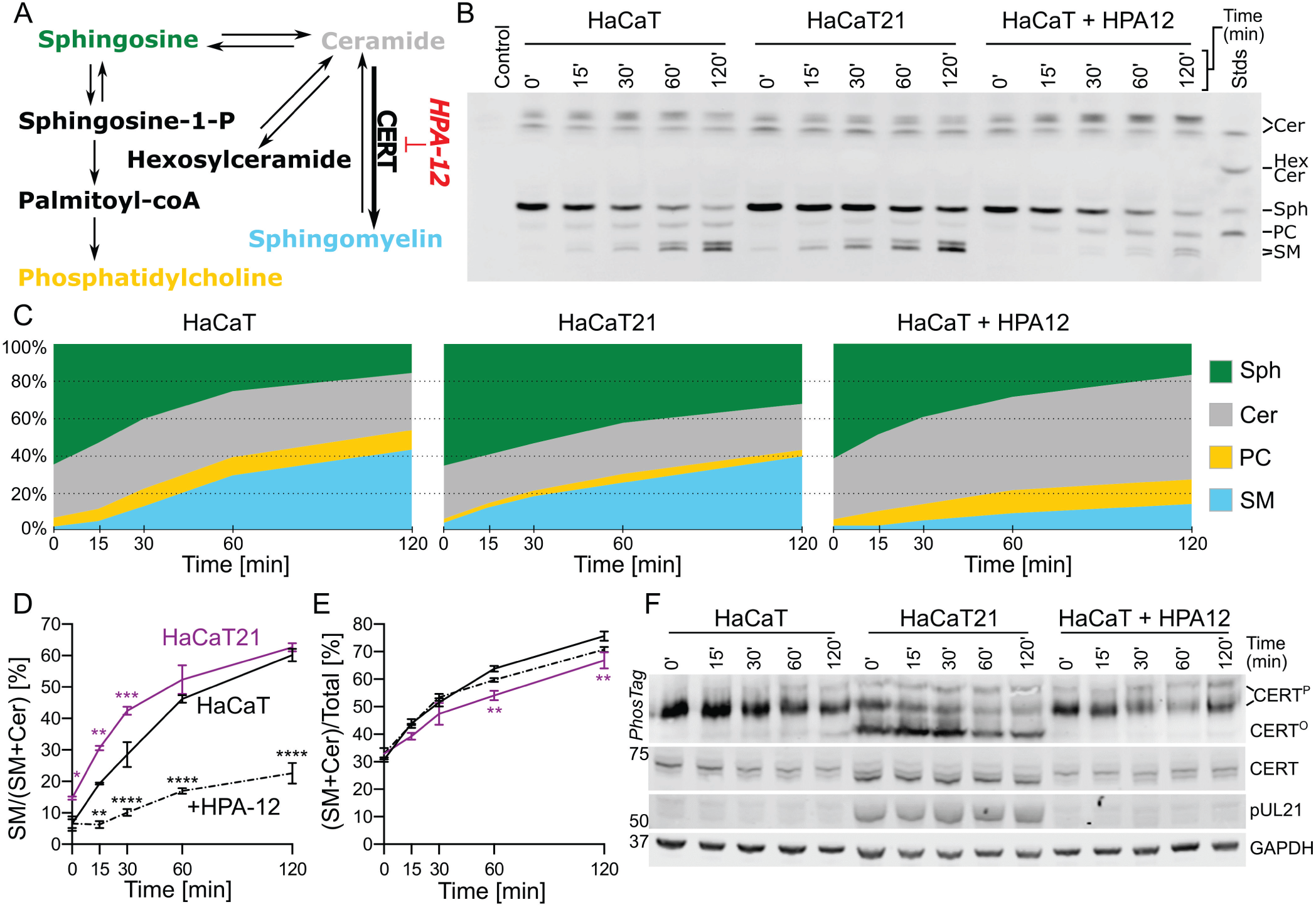
HSV-1 pUL21 alters the rate of ceramide to sphingomyelin conversion in cultured cells. (**A**) Simplified schematic diagram of sphingolipid biosynthetic pathways that lead from sphingosine (Sph) to sphingomyelin (SM) and glycosphingolipids (hexosylceramide; HexCer), or to phosphatidylcholine (PC) via a “salvage” pathway. Ceramide (Cer) to SM conversion is accelerated by the transport protein CERT, which is selectively inhibited by HPA-12. (**B**) Rate of Sph conversion to Cer, SM or PC was measured in HaCaT cells, untreated or treated with 1 µM HPA-12, and in HaCaT cells stably expressing pUL21 (HaCaT21). Cells were incubated with 5 μM ‘clickable’ alkyne-Sph (pulse) for 5 min and harvested for lipid extraction either immediately (0 min) or at the indicated times (chase). Extracted lipids were bioconjugated to 3-azido-7-hydroxycoumarin, separated by HPTLC and detected using UV light. Separated lipids were identified using clickable standards (Stds) and previous literature [42]. Data is from one representative experiment of two independent repeats. (**C**) Quantitation of the lipid intensities from (**B**) as determined by densitometry and represented as percentage fraction of total signal. (**D**) Rate of SM synthesis expressed as its fraction in the cumulative signal for SM and Cer. (**E**) The ratio of alkyne-Sph incorporated into either Cer or SM as a proportion of total alkyne-lipid signal, representing the influx of Sph into the SM biosynthetic pathway. For (**D**) and (**E**) the data represents two independent experiments (mean ± SEM). Data points are labelled if significantly different to parental HaCaT cells: **, p < 0.01; ***, p < 0.001; ****, p < 0.0001 (two-way ANOVA with Dunnett’s multiple comparisons test). (**F**) The re-solubilised proteins precipitated during lipid extraction were analysed by SDS-PAGE and immunoblotting using the antibodies listed. Where indicated, the gel was supplemented with PhosTag reagent to retard the migration of phosphorylated proteins, thus enhancing the separation of CERT that is hypo- (CERT^O^) or hyper-phosphorylated (CERT^P^). GAPDH serves as a loading control.

To monitor Sph metabolism, HaCaT cells, either parental or stably expressing pUL21 (HaCaT21), were incubated with alkyne-Sph for five minutes (pulse) and the rate of alkyne-Sph incorporation into the competing metabolic pathways was monitored for two hours (chase) by high performance thin layer chromatography (HPTLC) separation and detection of lipids conjugated to coumarin-azide via a ‘click’ reaction (Fig 1B and 1C). Alkyne-Sph was very efficiently converted to alkyne-Cer in both cell types, the levels of alkyne-Cer remaining relatively stable throughout the chase, and for both cell types synthesis of alkyne-PC and alkyne-SM was observed, but not synthesis of alkyne-HexCer. HaCaT21 cells exhibited significantly reduced rates of alkyne-Sph conversion and the level of alkyne-Cer is significantly lower (Fig S1). The rate of alkyne-PC synthesis was also reduced, although this reduction was not significant owing to the high inter-experiment variability of the alkyne-PC signal for the parental HaCaT cells. Taken together, these results indicated that lipid metabolism is altered in HaCaT cells constitutively expressing pUL21. However, it remained unclear whether these differences reflected adaptations of cellular lipid metabolism in response to constitutive pUL21 expression rather than a direct effect upon CERT activity. Since CERT-mediated non-vesicular transport of Cer from the ER to the TGN defines the rate of Cer-to-SM conversion [21–23], pUL21-directed CERT dephosphorylation (and thus activation) should increase the rate of SM synthesis. Consistent with this hypothesis, the abundance alkyne-SM as a fraction of total alkyne-Cer plus alkyne-SM was significantly higher in HaCaT21 versus HaCaT cells at early time points (0–30 min) during the chase (Fig 1D). By the 60 min chase time point the rate of alkyne-SM accumulation slows and the difference in relative abundance between cell lines was diminished, consistent with the alkyne-SM levels in both cell types approaching equilibrium. This increase in alkyne-SM accumulation as a fraction of alkyne-Cer plus alkyne-SM abundance was observed despite similar overall signal for alkyne-SM and alkyne-Cer in HaCaT21 cells when compared to parental cells at early time points (0–30 min), the abundance of these lipids being significantly lower in HaCaT21 cells at later time points (Fig 1E). Reduced overall levels of SM plus Cer in HaCaT21 cells likely results from adaptation to constitutive pUL21 expression (and thus constitutive CERT hyperactivation) via stimulation of feedback pathways that reduce Sph to Cer conversion and/or increased back-conversion of SM and/or Cer to Sph. Treatment of HaCaT cells with the highly-specific CERT inhibitor HPA-12 [44] confirmed that the rate of Cer-to-SM conversion was defined by CERT activity: SM synthesis was significantly decreased in HPA-12–treated cells when compared with untreated cells, and there was concomitant accumulation of alkyne-Cer in treated cells (Fig 1B–D and Fig S1). Immunoblot analysis of protein samples that were re-solubilized following their precipitation during lipid extraction (Fig 1F) confirmed that pUL21 was expressed and that dephosphorylated CERT (CERT^O^) predominates in HaCaT21 cells, in contrast to the parental and HPA-12 treated HaCaT cells.

pUL21 has multiple functions during infection, promoting the PP1-mediated dephosphorylation of not just CERT but of multiple different cellular and viral proteins [18]. It was therefore essential to identify a point mutant of pUL21 with impaired binding to CERT, but not to other cellular or viral partners, to dissect the functional significance of increased CERT activity during infection. Defining the interaction surfaces of transient protein complexes presents a considerable challenge [45], made more difficult in the case of pUL21 and CERT by their containing multiple domains and regions of intrinsic disorder [18]. The minimal region of CERT required for pUL21 binding was thus probed using pull-down experiments where pUL21-GST purified following bacterial expression (bait) was used to capture CERT or truncations thereof (preys) expressed by *in vitro* coupled transcription/translation in wheat germ extract (Fig 2A). The longer cytoplasmic isoform of CERT (CERT_L_), where expression of exon 11 extends the amino terminal region of the START domain by 26 amino acids (START_L_), was used since both CERT and CERT_L_ have been shown to possess ceramide transfer activity [23] and a contribution of these additional 26 amino acids to pUL21 binding could not be excluded. Full-length CERT_L_ was efficiently captured by pUL21-GST. The PH and START_L_ domains were also captured by pUL21-GST when expressed individually, albeit less efficiently, suggesting that the CERT_L_ possesses multiple pUL21- binding motifs. The CERT middle region, MR, was not captured by pUL21-GST and is thus likely to be dispensable for binding. A truncated CERT_L_ construct that retained the pUL21-binding PH and START_L_ domains, but lacked the majority of the highly flexible MR, was thus designed for use in subsequent biochemical and structural studies (miniCERT_L_) (Fig 2B). The extent of the MR retained to link the two domains was informed by the crystal structure of the CERT PH:START domain complex [46], defining a minimal distance of at least 55 Å between the PH domain C terminus and the START domain N terminus, and the desire to exclude the predicted coiled-coil region that may cause oligomerisation [26]. MR residues 132–350 were thus excluded from miniCERT_L_. The first 19 amino acids of the PH domain were also omitted from miniCERT_L_ as this region is predicted to be highly disordered and was excluded in previous structural studies [24, 47].

**Figure 2.**
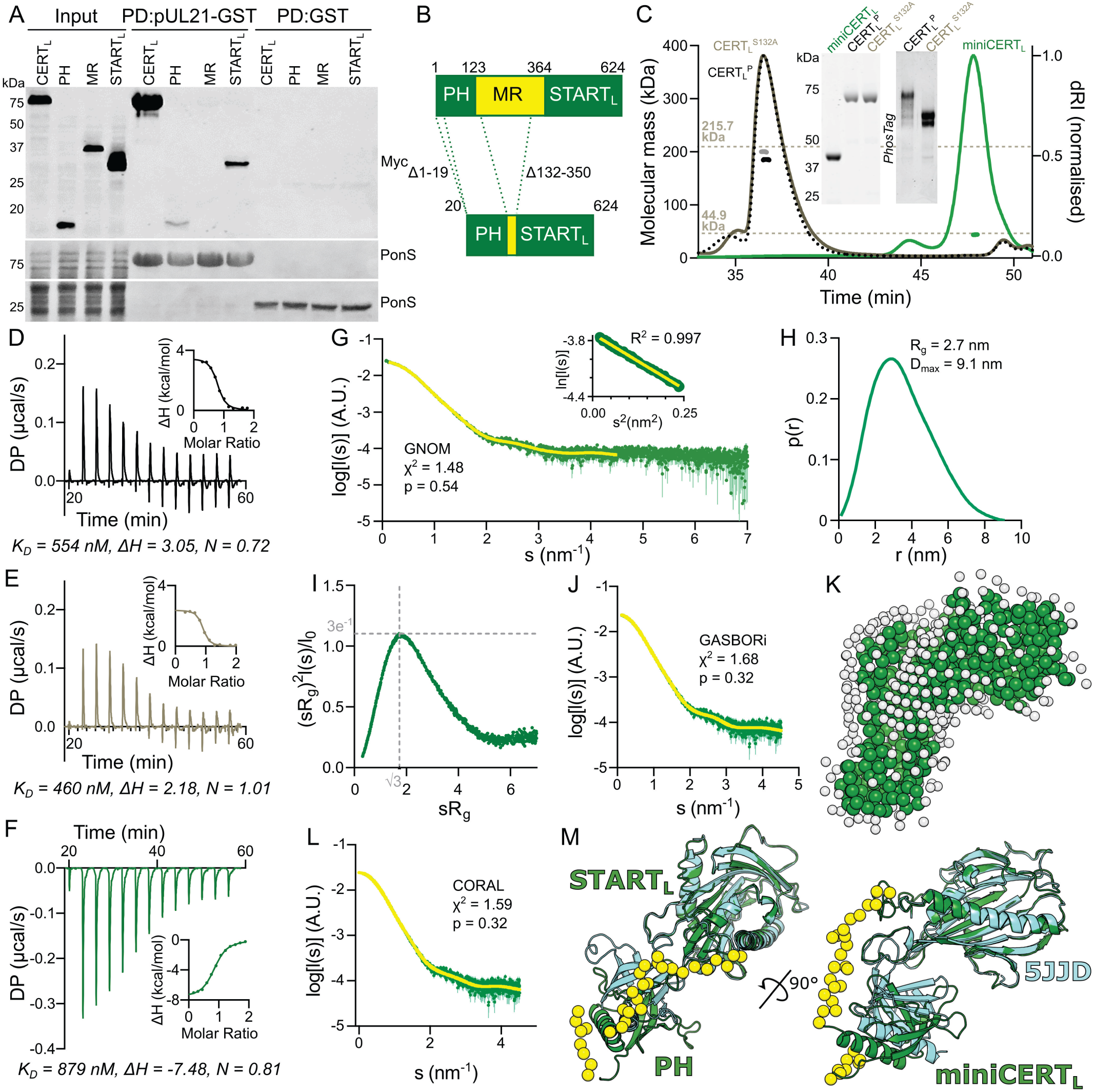
pUL21 binds to the PH and START_L_ domains of CERT_L_, and to a monomeric form of CERT_L_ comprising just the PH and START_L_ domains (miniCERT_L_). (**A**) Minimal binding elements of CERT_L_ were determined via pull down (PD) experiment using immobilised purified pUL21-GST or GST alone to capture myc-tagged full-length CERT_L_ or truncations thereof expressed via *in vitro* transcription/translation. Captured proteins were subjected to SDS-PAGE and immunoblotting using an anti-myc antibody. Ponceau S (PonS) staining of the immunoblot membrane before blocking shows equal and efficient capture of the bait proteins across the tested conditions. (**B**) Schematic representation of miniCERT_L_. Dotted lines indicate the regions of full-length CERT_L_ that were omitted. (**C**) SEC-MALS elution profiles (normalized differential refractive index, dRI) of StrepII-CERT_L_ (black dotted), StrepII-CERT_L_^S132A^ (grey solid) and H_6_-miniCERT_L_ (green solid). Weight-averaged molecular masses (coloured solid lines) are shown across the elution peaks. The expected molecular masses for trimeric StrepII-CERT_L_ and StrepII-CERT_L_^S132A^, and for monomeric H_6_-miniCERT_L_, are shown as dotted horizontal lines. Inset shows Coomassie-stained SDS-PAGE of the samples used for SEC-MALS (left), plus SDS-PAGE supplemented with PhosTag reagent (right) of the StrepII-CERT_L_ proteins. (**D–F**) Representative ITC titration curve of pUL21-H_6_ binding to (**D**) StrepII-CERT_L_, (**E**) StrepII-CERT_L_^S132A^, and (**F**) H_6_-miniCERT_L_. DP, differential power. Insets show normalized binding curves with integrated changes in enthalpy (ΔH) as a function of molar ratio. The affinity (*K_D_*), ΔH and stoichiometry (N) for the presented titrations are displayed below. (**G**) SAXS profile measured for H_6_-miniCERT_L_. The reciprocal- space fit of the *p*(*r*) profile to the SAXS data is shown as a yellow line. χ2, fit quality; p, Correlation Map (CorMap) probability of systematic deviations between the model fit and the scattering data [81]. Inset shows the Guinier plot (*sR_g_* < 1.3), which is linear as expected for an aggregate- and repulsion-free system. (**H**) The real-space distance distribution function, *p*(*r*), calculated from the SAXS profile. (**I**) Dimensionless Kratky plot of the SAXS data. Grey dotted lines indicate the expected maximum of the plot for a compact protein (*sR_g_* = √3, (*sR_g_*)^2^*I*(*s*)/*I*(0) = 3*e*^-1^) [91]. (**J**) Fit of an *ab initio* dummy-residue model calculated using GASBOR to the SAXS profile, and (**K**) representative GASBOR dummy-residue model. White spheres indicate modelled water beads of the hydration shell. (**L**) Fit to the SAXS profile of the pseudo-atomic model of H_6_-miniCERT_L_ obtained using CORAL. (**M**) CORAL pseudo-atomic model of H_6_-miniCERT_L_ (green ribbons) superimposed onto the crystal structure of the complex between the CERT PH and START domains (PDB ID 5JJD; cyan ribbons) [46] by aligning the START/START_L_ domains. Superposition is shown in orthogonal orientations where the linker regions or termini that were modelled by CORAL are depicted as yellow spheres.

Previous studies identified that the MR mediates CERT trimerisation [26, 48]. Size-exclusion chromatography with inline multi-angle light scattering (SEC-MALS) (Fig 2C) confirmed that hyperphosphorylated StrepII-tagged CERT_L_^P^, purified following expression in mammalian (Freestyle 293F) cells, has an observed mass (184.3 kDa) approaching that expected for a trimer (215.7 kDa). Similarly, a constitutively hypophosphorylated form of CERT_L_ where S132 is mutated to alanine, and thus cannot become phosphorylated to initiate the SRM phosphorylation cascade [28], had an observed mass (199.2 kDa) approaching that expected for a trimer. In contrast, H_6_-miniCERT_L_ purified following bacterial expression was predominantly monomeric and monodisperse (observed mass 43.9 kDa, expected monomeric mass 44.9 kDa), consistent with trimerization of CERT_L_ being driven by the MR and not being dependent upon CERT_L_ phosphorylation.

Isothermal titration calorimetry (ITC) demonstrated that pUL21 binds CERT_L_ with approximately micromolar affinity (Fig 2D,E and Table 1), the observed affinity not differing significantly between the hyperphosphorylated (CERT_L_^P^) and hypophosphorylated (CERT_L_^S132A^) forms of the protein. While CERT_L_^S132A^ forms a 1:1 complex with pUL21, the observed binding stoichiometry (N) was consistently lower for CERT_L_^P^ (0.74), consistent with a proportion of the pUL21 binding sites on CERT_L_ being sterically occluded when the protein is hyperphosphorylated. MiniCERT_L_ and pUL21 form an equimolar complex, binding with micromolar affinity similar to the full-length protein (Fig 2F; Table 1). However, the thermodynamics of binding differ significantly; whereas pUL21 binding to CERT_L_ is endothermic and entropically driven, binding to miniCERT_L_ is exothermic and enthalpically driven with minimal change in overall entropy (Table 1). This is consistent with CERT_L_ undergoing significant conformational rearrangement upon binding to pUL21, whereas the conformational changes to miniCERT_L_ (if any) upon pUL21 binding are likely to be much more modest.

**Table 1.** Thermodynamic properties of the pUL21:CERT_L_ interaction. . As quantitated by isothermal titration calorimetry (ITC). Experiments were performed *n* (replicates) times and mean ± SEM values are shown.

| Syringe | Cell | *K<sub>D</sub>* [nM] | Δ*H*<br>[kcal/mol] | Δ*G*<br>[kcal/mol] | -TΔ*S*<br>[kcal/mol] | N<br>(stoichiometry) | *n*<br>(replicates) |
| --- | --- | --- | --- | --- | --- | --- | --- |
| CERT<sub>L</sub><sup>P</sup> | pUL21 | 772.67 ±<br>158.18 | 3.89 ±<br>0.74 | -8.37 ± 0.11 | -12.27 ±<br>0.62 | 0.74 ± 0.01 | 3 |
| CERT<sub>L</sub><sup>S132A</sup> | pUL21 | 642.0 ±<br>182.0 | 2.385 ±<br>0.21 | -8.475 ±<br>0.18 | -10.85 ±<br>0.05 | 0.97 ± 0.04 | 2 |
| miniCERT<sub>L</sub> | pUL21 | 1042.0 ±<br>240.46 | -8.65 ± 0.28 | -8.21 ± 0.11 | 0.44 ±<br>0.35 | 0.89 ± 0.03 | 6 |
| miniCERT<sub>L</sub> | pUL21C | 4325.0 ±<br>1175.0 | -3.84 ± 0.02 | -7.35 ± 0.17 | -3.51 ±<br>0.18 | 0.97 ± 0.03 | 2 |
| miniCERT<sub>L</sub> | pUL21<sup>V382E</sup> | 8086.67 ±<br>538.53 | -7.20 ± 0.03 | -6.95 ± 0.04 | 0.26 ±<br>0.01 | 1.16 ± 0.08 | 3 |

Small-angle X-ray scattering (SAXS) in batch mode was used to probe the solution structure of H_6_- miniCERT_L_ (Fig 2G; Table S1). The frequency distribution of real-space distances (*p*(*r*) profile) is largely symmetric (Fig 2H) and the dimensionless Kratky plot (Fig 2I) has a bell-shaped peak at *sR_g_* = √3, consistent with miniCERT_L_ having a compact globular conformation in solution. *Ab initio* modelling of the H_6_-miniCERT_L_ scattering profile using GASBOR [49] yields a dummy-residue model (Fig 2J,K) that closely resembles the crystal structure of CERT START and PH domain complex [46]. A pseudo-atomic model of the H_6_-miniCERT_L_ solution structure, generated using the crystal structures of the CERT PH and START domains determined in isolation [24, 25] as rigid bodies, closely resembles the crystal structure of the CERT PH:START domain complex (Fig 2L,M). Collectively, the MALS, ITC and SAXS analysis demonstrates that miniCERT_L_ forms a compact monomeric protein with high affinity for pUL21, representing an excellent tool for structural studies.

Structural characterisation via X-ray crystallography and SAXS has revealed pUL21 to comprise two domains joined by a highly flexible linker [18,50,51], and immunoprecipitation experiments mapped CERT binding to the pUL21 C-terminal domain [18]. The C-terminal domain of pUL21 (pUL21C), spanning amino acids 275–535, was expressed with a hexahistidine tag and purified following bacterial expression. In contrast to a previous report [50] we did not observe co-purification of nucleic acid with H_6_-pUL21C, the purified protein having an A_260/280_ ratio of ∼0.6. SAXS analysis (Fig S2) confirmed that H_6_-pUL21C is monomeric and monodisperse, adopting a compact structure in solution that matches the previously determined crystal structure (χ^2^ = 0.99, CorMap p = 0.133) [50]. SEC analysis of a pre- formed complex between purified H_6_-miniCERT_L_ and a 1.34-fold molar excess of H_6_-pUL21C confirmed that the two proteins form a stable complex (Fig 3A), suitable for structural studies. ITC demonstrated that H_6_-pUL21C forms an equimolar complex with H_6_-miniCERT_L_ with a dissociation constant (*K_D_*) of 4.3 ± 1.2 μM (Fig 3B; Table 1). The approximately 4-fold reduction in miniCERT_L_ binding by pUL21C versus full-length pUL21 is consistent with previous immunoprecipitation results, where transfected pUL21C-GFP captured endogenous CERT slightly less efficiently than did pUL21-GFP in HEK293T cells [18]. These ITC experiments confirm that the C-terminal domain of pUL21 is the major determinant of CERT binding.

**Figure 3.**
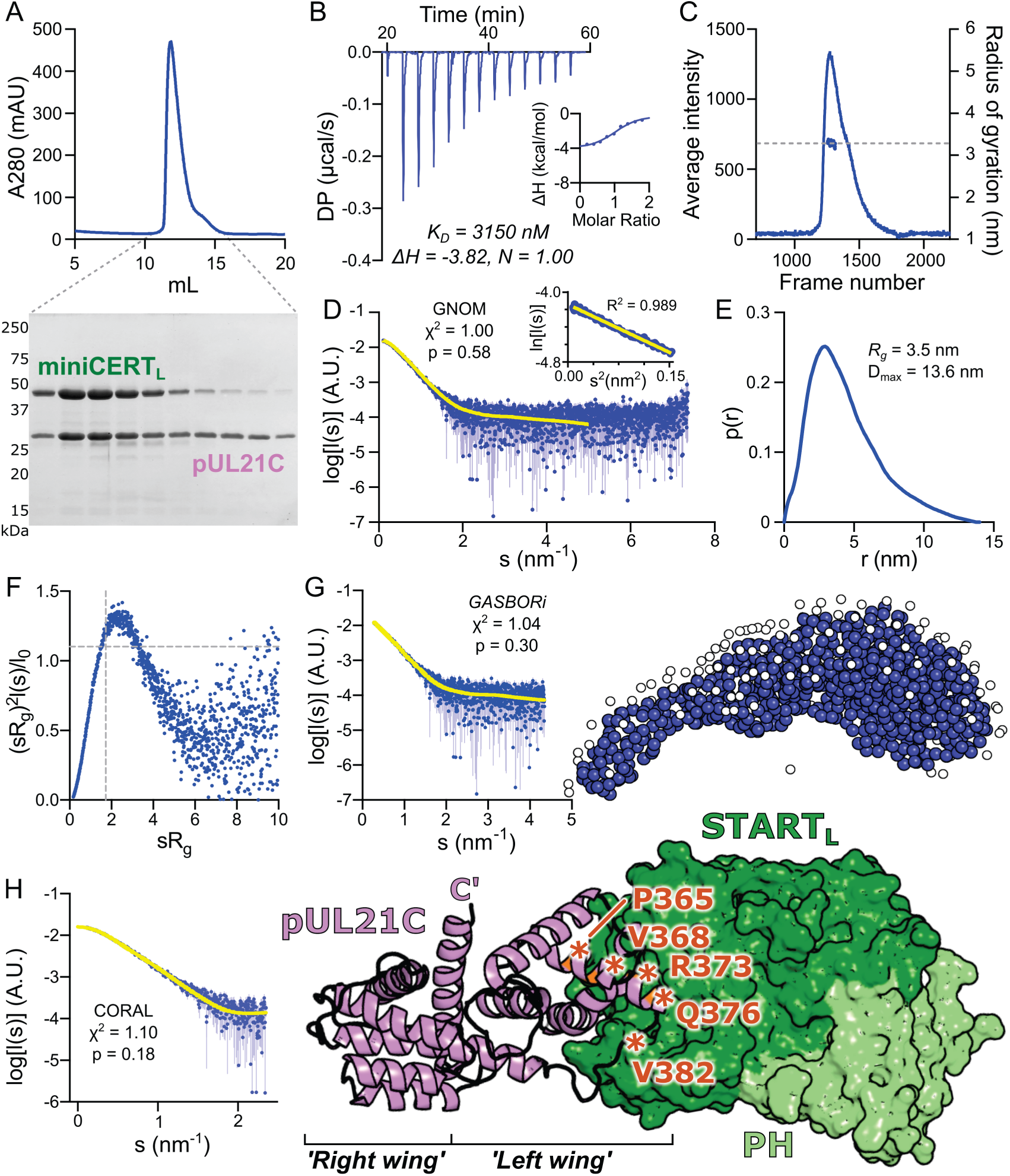
Solution structure of the H_6_-pUL21C:H_6_-miniCERT_L_ heterodimer. (**A**) SEC elution profile (A_280_) of the H_6_-pUL21C:H_6_-miniCERT_L_ complex, pre-formed in the presence of 1.32-fold molar excess of H_6_-pUL21C. Dotted lines indicate fractions that were collected and subjected to SDS-PAGE analysis with Coomassie staining, revealing co-elution of the two proteins. (**B**) Representative ITC titration curve of H_6_-pUL21C binding to H_6_-miniCERT_L_. DP, differential power. Inset shows normalized binding curve with integrated changes in enthalpy (ΔH) as a function of molar ratio. The ranges of the vertical axes are identical to Fig 2F. The affinity (*K_D_*), ΔH and stoichiometry (N) for the presented titration are displayed below. (**C**) SEC elution profile (partial integrated scattered X-ray intensity vs. data frame number) obtained during SEC-SAXS analysis of H_6_-pUL21C:H_6_-miniCERT_L_ complex. Dashed line indicates the calculated radius of gyration (*R_g_*) across the frames averaged for structural analyses. (**D**) Averaged SAXS profile of the H_6_-pUL21C:H_6_-miniCERT_L_ complex. The reciprocal-space fit of the *p*(*r*) profile to the SAXS data is shown as a yellow line. χ^2^, fit quality; p, Correlation Map (CorMap) probability of systematic deviations between the model fit and the scattering data [81]. Inset displays the Guinier plot (*sR_g_* < 1.3), which is linear as expected for an aggregate- and repulsion-free system. (**E**) Real- space distance distribution function, *p*(*r*), calculated from the SAXS profile. (**F**) Dimensionless Kratky plot of the SAXS data. Grey dotted lines indicate the expected maximum of the plot for a compact protein (*sR_g_* = √3, (*sR_g_*)^2^*I*(*s*)/*I*(0) = 3*e*^-1^). (**G**) *Ab initio* modelling of H_6_-miniCERT_L_:H_6_-pUL21C using GASBOR. Fit of the calculated scattering (yellow) to the SAXS profile is shown, as is a representative dummy-residue model (blue spheres) with modelled water beads of the hydration shell (white spheres). (**H**) Pseudo-atomic model of the H_6_-miniCERT_L_:H_6_-pUL21C complex generated using CORAL. The fit of the computed scattering (yellow) to the SAXS profile is shown. pUL21C is shown as a violet ribbon with C terminus labelled and miniCERT_L_ as a green molecular surface (PH, light green; START_L_, dark green). The left and right ‘wings’ of the dragonfly-like pUL21C fold [50] are labelled. For clarity, regions absent from the crystal structures that were modelled by CORAL are not displayed. Orange asterisks indicate pUL21C residues at the interface with miniCERT_L_ that were selected for further investigation. Additional pseudo-atomic models that also fit the scattering data are shown in Fig S3.

To probe the structural basis of the CERT recruitment by pUL21, a pre-formed complex of H_6_- miniCERT_L_ and H_6_-pUL21C was subjected to SEC with inline SAXS measurement (SEC-SAXS, Fig 3C). SAXS data were processed by averaging frames with a consistent calculated radius of gyration (*R_g_*) and then subtracting averaged buffer frames to yield the H_6_-miniCERT_L_:H_6_-pUL21C complex scattering profile (Fig 3D). The probable frequency of real-space distances (*p*(*r*) profile) of the complex is moderately asymmetric (Fig 3E), in contrast to the highly symmetric *p*(*r*) profiles of H_6_-pUL21C (Fig S2B) or H_6_-miniCERT (Fig 2H) alone, suggesting a less spherical particle, and the dimensions of the complex (*R_g_* 3.5 nm, D_max_ 13.6 nm) are substantially larger than for H_6_-pUL21C (*R_g_* 2.2 nm, D_max_ 8.5 nm) or H_6_-miniCERT (*R_g_* 2.7 nm, D_max_ 9.1 nm). The peak of the dimensionless Kratky plot is slightly higher, with its peak away from *sR_g_* = √3 (Fig 3F), suggesting some flexibility in the system granted either by modest dissociation of the complex or some mobility of the H_6_-miniCERT_L_ domains with respect to each other and H_6_-pUL21C. *Ab initio* modelling using GASBOR indicated an elongated molecule (Fig 3G). While initial pseudo-atomic models of the complex generated using a fixed conformation of H_6_-miniCERT_L_ did not fit the SAXS profile acceptably, allowing the PH and START_L_ domains freedom to move with respect to each other and to H_6_-pUL21C yielded three pseudo-atomic models with high-quality fits to the SAXS profile (Fig 3H and Fig S3). In all three top models miniCERT_L_ binds ‘end-on’ to the pUL21 molecule, forming an ellipsoidal ‘rugby ball’ like particle. In two of the top three models miniCERT_L_ binds the ‘left wing’ of the dragonfly-shaped pUL21C domain [50], whereas in the other it binds the ‘right wing’ (Fig S3). These models all have a similar overall shape, and thus all explain the SAXS scattering data well, but the relative orientations of H_6_-miniCERT_L_ and H_6_-pUL21C differ. All three top models were thus used to design specific pUL21 mutations that might disrupt (mini)CERT binding.

Mutations in pUL21C were designed to identify whether miniCERT_L_ binds the left or right wing of this domain. Amino acids in helix α4 or the subsequent loop of pUL21C were mutated to test binding to the left wing (Fig 3H, Fig S3), whereas amino acids in helices α5 and α9 were used to test binding to the right wing (Fig S3B). Immunoprecipitation experiments in transfected HEK293T cells that had been infected with HSV-1 lacking pUL21 expression (HSV-1 ΔpUL21) demonstrated that 4 of 5 substitution in the left wing of pUL21C (P365D, V368E, R373E and V382E) disrupted the ability of CERT to co-precipitate with pUL21-GFP (Fig 4A), whereas none in the pUL21C right wing disrupted CERT co-precipitation (Fig S3C). These results are consistent with CERT binding the left wing of pUL21C. The proximity of the pUL21C amino terminus to miniCERT_L_ in these models is also consistent with observations that CERT binding is lost when pUL21C has a bulky N-terminal GFP tag, but retained when the GFP tag is C-terminal [18].

**Figure 4.**
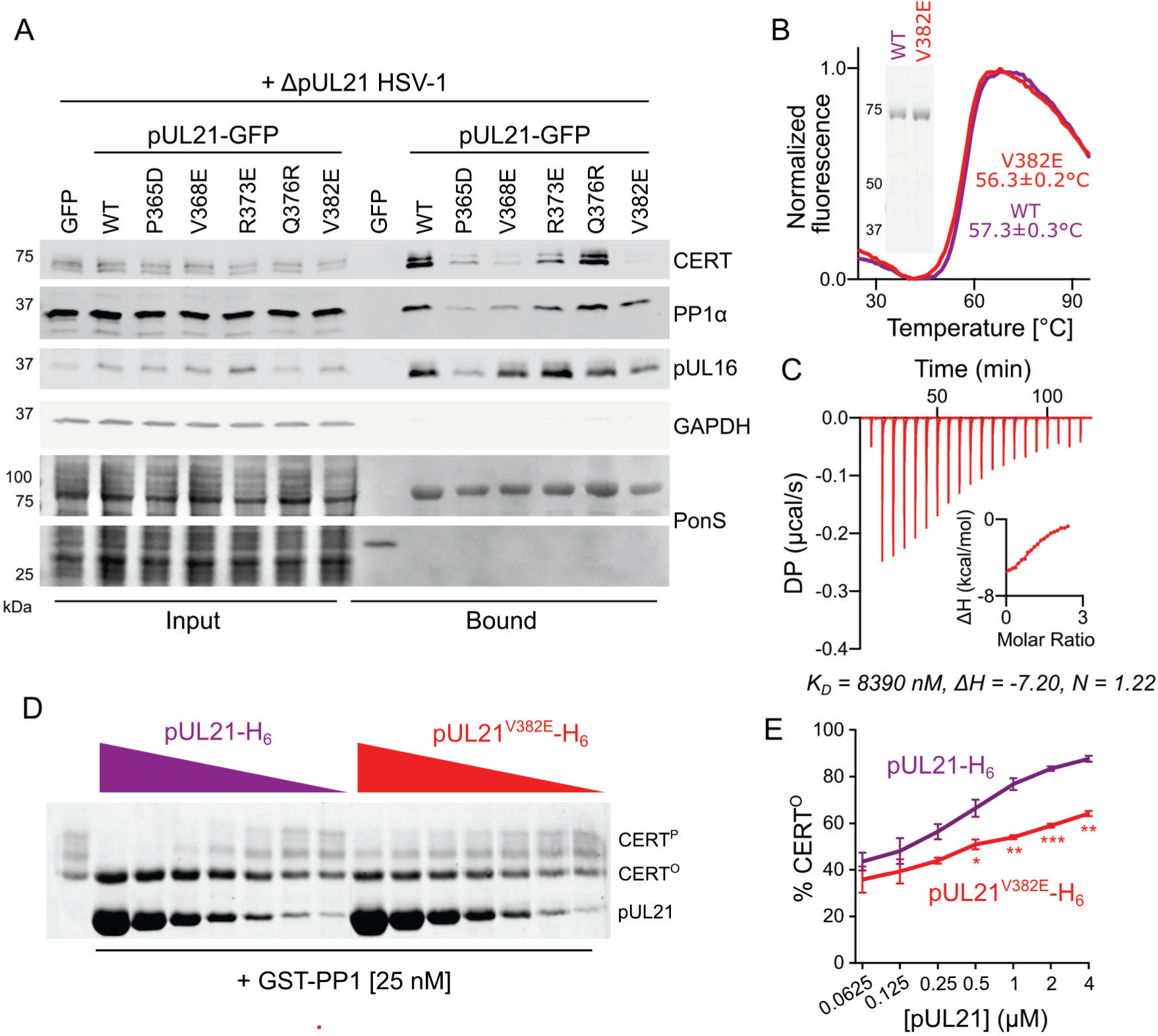
Identification and validation of a pUL21 point mutant with disrupted binding to CERT. (**A**) Immunoblot following immunoprecipitation from HEK293T cells transfected with plasmids encoding GFP-tagged pUL21, either wild-type (WT) or with amino acid substitutions at the putative CERT_L_ binding interface (Fig 3H), or encoding GFP alone. At 24 hours post-transfection cells were infected with ΔpUL21 HSV-1 (MOI = 5) and at 16 hours post-infection cells were lysed, tagged proteins were captured using GFP affinity resin, and the bound proteins were subjected to SDS-PAGE and immunoblotting using the antibodies listed. Ponceau S (PonS) staining of the nitrocellulose membrane before blocking is shown to confirm efficient capture of GFP-tagged proteins. (**B**) Differential scanning fluorimetry of WT (purple) and V382E (red) pUL21-H_6_. Representative curves are shown and melting temperature (T_m_) is mean ± SEM of three technical replicates. Inset shows Coomassie-stained SDS-PAGE of the purified proteins. (**C**) Representative ITC titration curve of pUL21^V382E^-H_6_ binding to H_6_-miniCERT_L_. DP, differential power. Inset shows normalized binding curve with integrated changes in enthalpy (ΔH) as a function of molar ratio. The ranges of the vertical axes are identical to Fig 2F. The affinity (*K_D_*), ΔH and stoichiometry (N) for the presented titration are displayed below. (**D**) *In vitro* dephosphorylation assay using all-purified proteins. 0.5 μM CERT^P^ was incubated with varying concentrations of pUL21-H_6_ WT or V382E (two-fold dilution from 4 μM to 62.5 nM) in the presence of 25 nM GST-PP1 for 30 min at 30°C. Proteins were resolved using SDS-PAGE supplemented with PhosTag reagent to enhance separation of CERT that is hyper- (CERT^P^) or hypo-phosphorylated (CERT^O^), and protein bands were visualised using Coomassie. Images are representative of two independent experiments. (**E**) Quantitation of concentration-dependent pUL21-mediated stimulation of CERT dephosphorylation as determined by densitometry. Ratio of CERT^O^ to total CERT (CERT^O^+CERT^P^) for two independent experiments is shown (mean ± SEM). Data points are labelled if significantly different: *, p < 0.05; **, p < 0.01; ***, p < 0.001 (two-way ANOVA with Sidak’s multiple comparisons test)

Of the pUL21 substitutions that disrupted CERT binding (Fig 4A), V382E appeared to cause the largest decrease in CERT binding while maintaining the ability of pUL21 to co-precipitate its other known binding partners, PP1 [18] and pUL16 [10]. H_6_-pUL21^V382E^ was purified following bacterial expression. Differential scanning fluorimetry (a.k.a. Thermofluor) showed pUL21^V382E^ to be well folded as its thermal stability is similar to wild-type H_6_-pUL21 (Fig 4B). ITC analysis demonstrated that H_6_-pUL21^V382E^ has approximately 8-fold reduced binding affinity for H_6_-miniCERT_L_ when compared to wild-type pUL21 (Fig 4C; Table 1). The effect of the V382E substitution upon the ability of pUL21 to promote CERT dephosphorylation, converting CERT^P^ to CERT^O^, was probed using an *in vitro* dephosphorylation assay with all-purified reagents (Fig 4D and E). The dose-dependent acceleration of GST-PP1 mediated CERT dephosphorylation is significantly greater for wild-type pUL21 than pUL21^V382E^ (p = 0.040; two-way ANOVA with Sidak’s multiple comparison test), consistent with the GST-PP1:pUL21^V382E^ complex having lower affinity for the substrate CERT^P^ (EC50[pUL21] = 0.491 ± 0.123 µM, EC50[pUL21^V382E^] = 3.329 ± 0.7874 µM; three parameter dose-response curve, n = 2 independent experiments).

To probe the ability of pUL21^V382E^ to stimulate CERT dephosphorylation during infection, a mutant strain of HSV-1 encoding pUL21^V382E^ was generated using two-step Red recombination [52]. Dephosphorylated CERT (CERT^O^) is significantly less abundant in HaCaT cells infected with HSV-1 expressing pUL21^V382E^ or lacking pUL21 expression (ΔpUL21) when compared to wild-type HSV-1 infected cells (Fig 5A,B). In addition to CERT, pUL21 expression reduces the phosphorylation of multiple substrates of the viral kinase pUS3 in HSV-1 infected cells [18], the phosphorylated forms of these substrates being detectable using an antibody that recognises phosphorylated substrates of the cellular kinase Akt [53]. While infection with ΔpUL21 HSV-1 causes a dramatic increase in the abundance of multiple phosphorylated pUS3 substrates, the abundance of these phosphoforms is indistinguishable between wild-type and pUL21^V382E^ HSV-1 (Fig 5A). This confirms that the V382E substitution specifically disrupts CERT dephosphorylation, rather than generally inhibiting the ability of pUL21^V382E^ to recruit PP1 to substrates. Similar changes in CERT dephosphorylation, but not in the dephosphorylation of other pUS3 substrates, are observed when Vero cells are infected with pUL21^V382E^ HSV-1 (Fig S4A,B). Immunocytochemistry confirms that both wild-type and V382E pUL21 have the similar subcellular localisation, being observed predominantly at the nuclear rim of infected Vero cells (Fig 5C).

**Figure 5.**
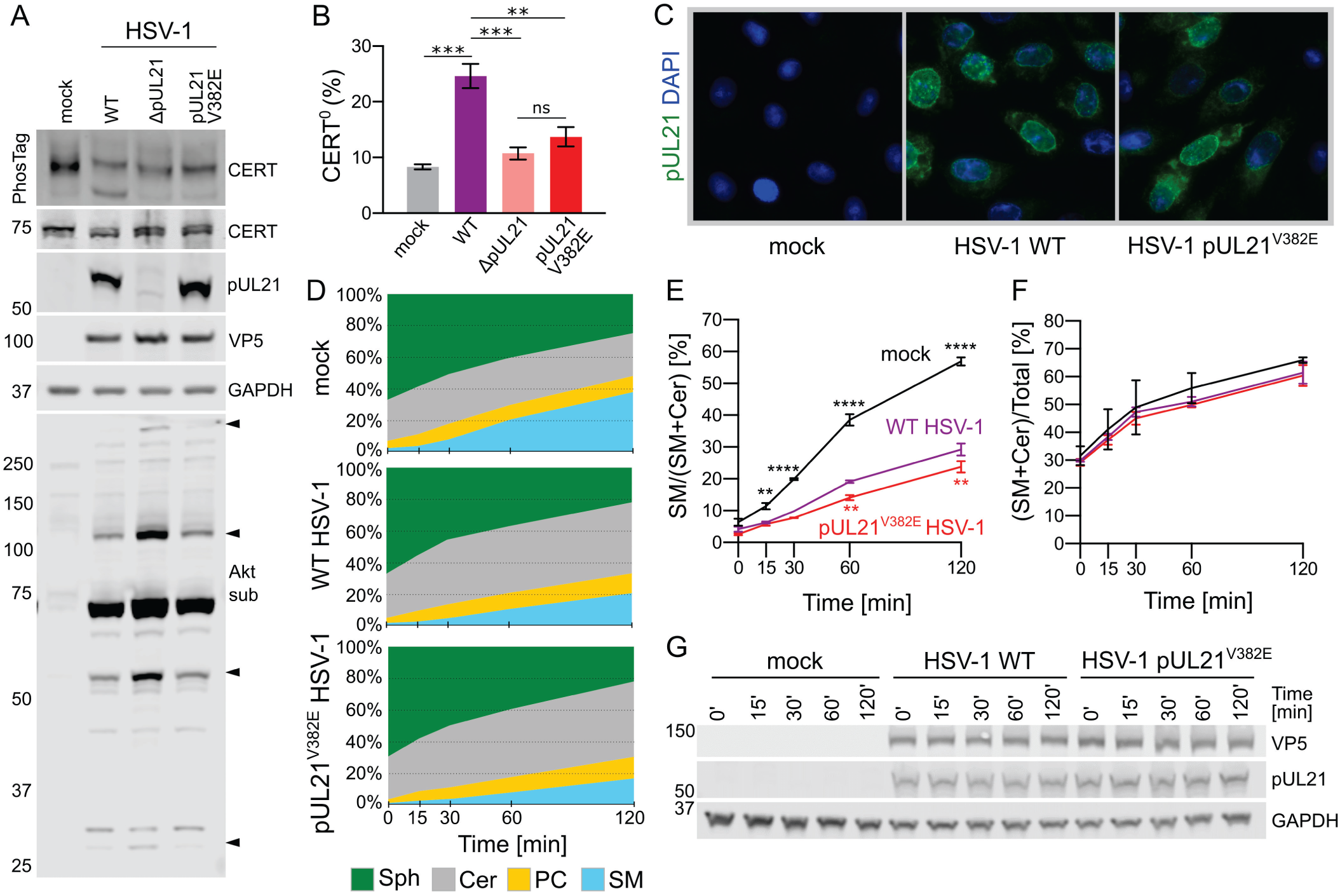
Mutating the CERT-binding interface of pUL21 inhibits CERT dephosphorylation and reduces the rate of sphingomyelin synthesis in infected cells. (**A**) HaCaT cells were infected at MOI = 5 with wild-type (WT) HSV-1, HSV-1 lacking pUL21 expression (ΔpUL21), or a pUL21 point mutant virus (pUL21^V382E^). Lysates were harvested at 16 hpi in the presence of phosphatase inhibitors and subjected to SDS-PAGE plus immunoblotting using the antibodies listed. Where indicated, the gel was supplemented with PhosTag reagent to enhance separation of CERT phosphoforms. The antibody recognising phosphorylated Akt substrates (Akt sub) illustrates activity of the HSV-1 kinase pUS3, several substrates of which are dephosphorylated in a pUL21-dependent manner (arrowheads) [18]. VP5, infection control. (**B**) Quantitation of the CERT dephosphorylation level (ratio of CERT^O^ to total CERT) in cells infected with WT or mutant HSV-1, as determined by densitometry. Results are presented as mean ± SEM from three independent experiments. One-way ANOVA with Dunnett’s multiple comparisons test was used for the statistical analysis (ns, non-significant; **, p < 0.01; ***, p < 0.001). (**C**) Vero cells were infected at MOI = 1, fixed 14 hpi and stained with DAPI (blue) plus an antibody recognising pUL21 (green). (**D**) Pulse-chase experiment to measure the rate of Sph conversion to Cer, SM and PC. HaCaT cells were infected with WT or pUL21^V382E^ HSV-1 at MOI = 5, or mock infected. Cells were incubated with 5 μM alkyne-Sph (pulse) at 14 hpi for 5 min and harvested for lipid extraction either immediately (0 min) or at the indicated times (chase). Extracted lipids were bioconjugated to 3-azido-7-hydroxycoumarin using click chemistry, separated by HPTLC, visualised using UV light and relative lipid abundances were quantitated by densitometry. Data is from one representative experiment of two independent repeats. (**E**) Rate of SM synthesis expressed as its fraction in the cumulative signal for SM and Cer. (**F**) The proportion of click-Sph incorporated into either Cer or SM, representing the temporal influx of Sph into the SM biosynthesis pathway. For (**E**) and (**F**) the data represents two independent experiments (mean ± SEM). Data points are labelled if significantly different to WT HSV-1: **, p < 0.01; ****, p < 0.0001 (two-way ANOVA with Dunnett’s multiple comparisons test). (**G**) The re-solubilised proteins precipitated during lipid extraction were analysed by SDS-PAGE and immunoblotting using the antibodies listed.

Metabolic labelling was used to monitor the impact of pUL21-mediated CERT dephosphorylation on sphingolipid biogenesis during infection. A pulse-chase experiment was performed where HaCaT cells infected with wild-type or pUL21^V382E^ HSV-1, or mock infected, were incubated for 5 min with alkyne- Sph at 14 hours post-infection (hpi) and its metabolites were monitored for two hours. Cer accumulates and the rate of SM synthesis is significantly decreased in cells infected with wild-type and pUL21^V382E^ HSV-1 when compared to uninfected cells (Fig 5D). Although a substantial decrease in the rate of SM synthesis is seen for both wild-type and mutant HSV-1 infection, the defect is significantly larger in cells infected with HSV-1 pUL21^V382E^ (Fig 5E). The overall abundance of SM plus Cer is similar between infected and uninfected cells, suggesting that Cer to SM conversion is specifically impaired rather than influx of click-Sph into the SM biogenesis pathway being defective (Fig 5F). Taken together, this suggests that pUL21-mediated activation of CERT accelerates Cer to SM conversion during infection, albeit from a much lower base than in uninfected cells.

Having confirmed that pUL21^V382E^ HSV-1 specifically lacks the ability to stimulate CERT dephosphorylation, and that HSV-1 encoding pUL21^V382E^ has a reduced rate of Cer to SM conversion, the impact of this deficit upon virus replication and spread in cultured cells was assessed. Wild-type and pUL21^V382E^ HSV-1 form similar sized plaques on HaCaT and Vero cells (Fig 6A), suggesting that CERT dephosphorylation is dispensable for efficient viral cell-to-cell spread. A single-step growth curve, where Vero and HaCaT cells are infected at a high multiplicity of infection (MOI) and the production of infectious progeny is monitored over time, was used to compare the replication of wild type and pUL21^V382E^ HSV-1 (Fig 6B). Both viruses produce similar abundance of infectious progeny by 24 hours post infection. In two biologically independent experiments performed for each cell type the kinetics of virus replication was consistently accelerated, with higher titres of pUL21^V382E^ HSV-1 being observed between 6–12 hours post-infection, but the difference in growth rate is not statistically significant for either cell line (two-way ANOVA with Sidak’s multiple comparisons test).

**Figure 6.**
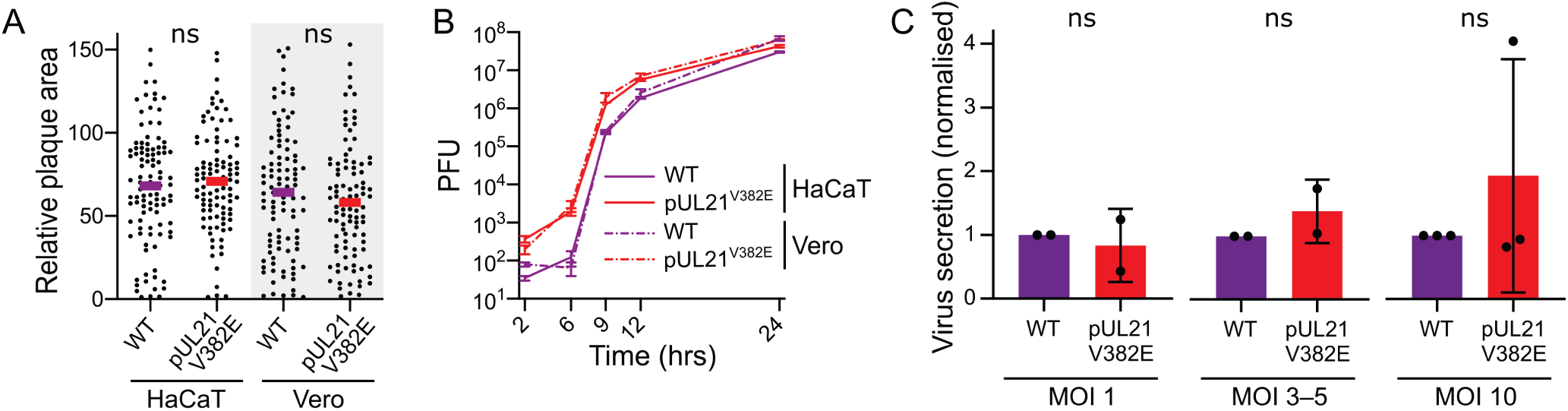
pUL21-mediated dephosphorylation of CERT accelerates virus replication but does not enhance virus secretion or cell-to-cell spread. (**A**) Monolayers of HaCaT or Vero cells were infected with 100 plaque forming units of WT or pUL21^V382E^ HSV-1. Following infection, cells were overlaid with medium containing 0.6% caboxymethyl cellulose and incubated for 48 hours, then fixed and immunostained with chromogenic detection. Relative plaque areas (pixels) were measured using Fiji [88, 89]. Bars represent mean plaque sizes, which were compared using unpaired t-test (n=100; ns, non-significant). (**B**) Single-step (high MOI) growth curves of WT and pUL21^V382E^ HSV-1. Monolayers of HaCaT (continuous line) or Vero (dotted line) cells were infected (MOI = 5) with the viruses shown. Samples were harvested at the indicated times and titres determined by plaque assay using Vero cells. Data are presented as mean values ± SEM of technical duplicates from one representative experiment. Difference in replication kinetics across two biological replicates is not statistically significant (two-way ANOVA with Sidak’s multiple comparisons test). (**C**) Virus release into the culture supernatant from HaCaT cells infected with WT or pUL21^V382E^ HSV-1 at various MOIs. Samples were harvested at 12 hpi and virus infectivity in the cells versus the culture medium was measured by titration on Vero cells. The fold change in secretion of infectivity into the culture medium for pUL21^V382E^ versus WT HSV-1 is shown as mean values ± SEM of two (MOI = 1 or 3–5) or three (MOI = 10) independent experiments. For MOI 3–5, the data represent one independent experiment performed at MOI = 3 and one at MOI = 5. For each MOI, the extent of virus secretion was compared using an unpaired t-test (ns, non-significant).

While HSV-1 preferentially remains cell-attached, spreading via direct cell:cell contacts, the cell-free secretion of virions from infected cells is altered when CERT is depleted or overexpressed [54]. Cell-free release of pUL21^V382E^ HSV-1 was determined by quantitating the amount of infectivity secreted into the medium as a percentage of overall infectivity (secreted plus cell-associated), normalised for each independent replicate to the secretion of a wild-type virus in control experiments performed at the same time. HaCaT cells infected for 12 h with various MOI of pUL21^V382E^ HSV-1 did not release significantly more or less infectivity when compared to WT HSV-1 infection (Fig 6C). Similarly, the cell-free release of infectivity at 12 hpi did not differ between Vero cells infected with pUL21^V382E^ or WT HSV-1 (Fig S4C).

## Discussion

HSV-1 extensively remodels the proteome of infected cells, altering both the abundance [3] and post-translational modification status [2,18,55] of multiple cellular proteins. We know much less about HSV-1 mediated changes to the cellular lipidome, although previous studies have identified that HSV-1 infection changes phosphoinositide levels [56] and increases the rate of *de novo* phospholipid synthesis [57]. By combining biochemical and structural studies with cell-based models of infection we show here that pUL21 accelerates the conversion of Cer to SM in infected cells by promoting dephosphorylation and activation of the Cer transport protein CERT (Fig 5). This is the first observation of a viral protein directly binding CERT and altering its activity. While pUL21-mediated CERT activation causes an apparent modest acceleration in the rate of virus replication, this effect is not statistically significant and CERT activation does not contribute to replication or cell-to-cell spread of HSV-1 in cultured keratinocytes or epithelial cells (Fig 6).

Our previous work showed that pUL21 is a phosphatase adaptor with multiple different targets [18]. Mutation of the pUL21 TROPPO motif, required for PP1 binding and thus stimulation of dephosphorylation, dramatically reduced the replication and spread of HSV-1. While the TROPPO motif was identified via its conservation across alphaherpesvirus sequences, the molecular basis of CERT recruitment remained unknown. Biochemical mapping and SAXS structural characterisation allowed us to identify a specific mutation, V382E, that decreases the affinity of pUL21 for miniCERT_L_ by approximately 8-fold (Fig 4C). While this reduction in affinity is moderate when assayed in dilute biochemical solution, it is sufficient to disrupt co-precipitation of CERT from transfected cells (Fig 4A), it abolishes pUL21-mediated dephosphorylation of CERT during infection (Fig 5A,B), and it significantly increases the rate of Cer to SM conversion (Fig 5E). These results are consistent with other binding partners competing to bind CERT and/or pUL21 within the context of infected cells, these competing interactions amplifying the effect of reduced pUL21:CERT binding affinities. The V382E mutation does not prevent pUL21 binding to pUL16 (Fig 4A) nor its ability to stimulate dephosphorylation of other targets (Fig 5A), confirming that pUL21 binds CERT via a molecular surface that is distinct from the binding site(s) of other targets. The wild-type levels of virus replication and cell-to-cell spread observed for pUL21^V382E^ HSV-1 are consistent with our previous *in vitro* evolution studies of HSV-1 mutants where the pUL21 TROPPO motif was mutated [18]. Adaptation of the virus to the loss of pUL21 PP1 binding, via suppressor mutations that reduce the activity of the kinase pUS3, restored virus replication and spread without restoring enhanced CERT dephosphorylation [18]. We therefore conclude CERT is not a critical substrate of pUL21 for virus replication and spread in cultured fibroblasts and keratinocytes. However, we note that HSV-1 is a neurotropic virus and that sphingolipids like SM are highly enriched in neurons, where they are critical for correct neuronal development and function [31]. Studying the role of pUL21-mediated CERT dephosphorylation in neurons thus remains an important question for future investigation.

Studies of cellular lipid metabolism are complicated by robust cellular feedback mechanisms [58] and challenges in accurately detecting changes in lipid abundance [59]. Here, we successfully employed lipid labelling, click chemistry and HPTLC to monitor the kinetics of sphingosine metabolism, revealing that HSV-1 pUL21 significantly increases the rate of CERT-mediated Cer to SM conversion both outside (Fig 1) and within (Fig 5) the context of infection. Furthermore, we observe a highly significant increase in the abundance of labelled Cer when cells are infected with HSV-1, this increase being greater when pUL21-mediated CERT activation has been abolished. The dramatic change in the rate of Cer to SM conversion is most likely explained by the known propensity of HSV-1 to promote dispersal of the Golgi and TGN [4, 60]. Such dispersal would alter ER:TGN contact sites, where CERT-mediated transport enables Cer to SM conversion by TGN-resident sphingomyelin synthase [61].

HSV-1 utilizes the PKD-mediated trafficking from the TGN to the plasma membrane [62]. Disruption of PKD-mediated trafficking has been shown to alter virus secretion, with siRNA-mediated depletion of CERT increasing the secretion of extracellular (cell-free) HSV-1 particles and CERT overexpression reducing virus secretion [54]. We do not observe changes in the secretion of pUL21^V382E^ HSV-1, which lacks the ability to dephosphorylate CERT and thus activate CERT-mediated Cer transport. While these results may appear at first glance to be contradictory, it should be noted that in our experiments the cells contained wild-type levels of CERT and cells infected with either ΔpUL21 or pUL21^V382E^ HSV-1 have similar levels of active (CERT^P^) and inactive (CERT^O^) protein compared to uninfected controls (Fig 5A). In contrast, siRNA depletion would lead to a complete absence of CERT and of CERT-mediated lipid transfer. Furthermore, the previous study demonstrated that treatment of infected cells with HPA-12 did not stimulate virus particle secretion [54]. This insensitivity of HSV-1 secretion to CERT pharmacological inhibition, combined with our observation that pUL21-mediated CERT activation does not alter HSV-1 secretion, strongly suggests that CERT catalytic activity is not directly linked to the regulation of virus secretion.

Our observation of labelled Cer accumulation during HSV-1 infection (Fig 5) is consistent with an earlier report that HSV-1 infection causes an approximately 2-fold increase in Cer abundance within BHK-21 cells [34]. Cer has many distinctive physical properties that set it apart from other membrane lipids: it has negative intrinsic curvature, it increases the order of phospholipids in membranes, and it makes biological membranes more permeable to even large solutes such as proteins [63]. Nascent HSV-1 particles must traverse multiple biological membranes during virus assembly, most notably during capsid egress from the nucleus. Being too large to egress via nuclear pores, capsids leave by budding into and then out of the perinuclear space, the former step being catalysed by the herpesvirus nuclear egress complex (NEC) [64]. A recent study identified that the NEC induces lipid ordering to generate the negative curvature required for capsid budding [65]. It is tempting to speculate that the accelerated replication kinetics of pUL21^V382E^ HSV-1 arises, at least in part, from an increased rate of nuclear capsid egress owing to increased Cer abundance. This effect would be distinct from the previously demonstrated role of pUL21 in promoting nuclear egress via regulating the phosphorylation of NEC components [18, 20]. Increased Cer abundance could also stimulate secondary envelopment of virus particles at post-Golgi membranes. However, in addition to potentially promoting virus assembly, accumulation of Cer can lead to caspase 3 activation and apoptosis via increased mitochondrial outer membrane permeabilization [66], and HSV-1 is known to encode proteins that defend against ceramide-induced apoptosis [67]. It is therefore possible that pUL21-stimulated acceleration of Cer to SM conversion acts to limit the pro-apoptotic activity of Cer, although confirmation of this requires further study.

In summary, we demonstrated that pUL21 dephosphorylates and activates the cellular lipid transport protein CERT, stimulating conversion of Cer to SM. Characterising the solution structure in complex with the PH and START domains of CERT allowed us to identify a single amino acid mutation of pUL21 that disrupts CERT dephosphorylation in infected cells. HSV-1 encoding this pUL21 mutant had similar replication kinetics, virus yields and plaque sizes, confirming that dephosphorylation of other cellular and/or viral targets underpins the important role of pUL21 in HSV-1 replication and spread. The functional rationale for pUL21-mediated modulation of CERT activity by HSV-1 remains elusive, but we have defined the molecular tools that will allow its dissection in other cell types and/or animal models of infection.

## Materials and methods

### Plasmids

The StrepII-CERT transient mammalian expression construct was described previously [18]. For generation of stable cells, a synthetic gene encoding human CERT_L_ (UniProt Q9Y5P4-1) was cloned into a modified version of plasmid PB-T [68] that encodes an N-terminal StrepII tag and a woodchuck hepatitis virus posttranscriptional regulatory element [69] in the 3′ untranslated region. The S132A substitution was introduced into this vector using QuikChange mutagenesis (Agilent) according to the manufacturer’s instructions. CERT_L_ destined for expression using *in vitro* transcription/translation system from wheat germ extract was cloned into the plasmid pF3A WG BYDV (Promega) with an N-terminal myc epitope tag. The truncations of CERT_L_ were generated by inverse PCR (MR, residues 123–364; START_L_, residues 365–624) or by introduction of a stop codon (PH, residues 1–122) using QuikChange mutagenesis. For purification following bacterial expression, miniCERT_L_ (residues 20–131 plus 351–624) was cloned into pOPTH [70], encoding a N-terminal MAH_6_ tag. The generation of pUL21- H_6_ and pUL21-GFP were described previously [18] and single amino acid substitutions were introduced by QuikChange mutagenesis. pUL21-H_6_ was subcloned into pOPTH and inverse PCR was used to generate H_6_-pUL21C, encoding pUL21 amino acids 275–535. A plasmid (UK622) encoding mouse PP1γ (UniProt P63087) with an N-terminal GST tag [71] was a kind gift from David Ron (Cambridge Institute for Medical Research). To generate pUL21^V382E^ HSV-1, pEPkan-S containing an I-SceI/KanR selection cassette was used [52].

### Mammalian cell culture

Mycoplasma-free spontaneously immortalised human keratinocyte (HaCaT) cells [72], HaCaT cells stably expressing pUL21 (HaCaT21) [18], African green monkey kidney (Vero) cells (ATCC #CRL- 1586) and human embryonic kidney (HEK)293T cells (ATCC #CRL-3216) were maintained in Dulbecco’s Modified Eagle Medium with high glucose (Merck), supplemented with 10% (v/v) heat-inactivated foetal calf serum (FCS) and 2 mM L-glutamine (complete DMEM) in a humidified 5% CO_2_ atmosphere at 37°C. For protein purification, Freestyle 293F suspension cells (ThermoFisher) were grown in Freestyle 293F medium (Gibco) on a shaking platform (125 rpm) in a humidified 8% CO_2_ atmosphere at 37°C.

Doxycycline-inducible stably transfected Freestyle 293F cells expressing StrepII-CERT_L_ and StrepII- CERT_L_^S132A^ were generated using a piggyBac transposon-based system [68]. A 30 mL suspension culture of Freestyle 293F cells at 1×10^6^ cells/mL were transfected with a 5:1:1 mass ratio of PB-T- CERT_L_^(S132A)^:PB-RN:PBase (35 μg total DNA) using Freestyle MAX transfection reagent (Invitrogen) as per the manufacturer’s instructions. After two days the cells were transferred to fresh media supplemented with 500 μg/mL Geneticin (Gibco) and the drug selection was continued for two weeks with media replenishment every three days.

### GFP affinity capture

Monolayers of HEK293T cells grown in 9 cm dishes were transfected with TransIT-LT1 (Mirus) using 7.7 μg of pEGFP-N1 (for GFP alone), pUL21-GFP, or point mutants thereof, following the manufacturer’s instructions. At 24 hours post-transfection, the cells were harvested by scraping into the medium, pelleted (220 g, 5 min, 4°C), washed three times with cold phosphate buffered saline (PBS) and lysed at 4°C in 1 mL lysis buffer (10 mM Tris pH 7.5, 150 mM NaCl, 0.5 mM EDTA, 0.5% NP-40, 1:100 diluted EDTA-free protease inhibitor cocktail [Merck]) for 45 min before clarification (21,000×g, 10 min, 4°C). After immunoprecipitation with GFP-Trap beads (ChromoTek) performed in accordance with the manufacturer’s protocol, the samples were eluted by incubation at 95°C for 5 min in 45 μL 2× SDS-PAGE loading buffer. Input and bound samples were separated by SDS-PAGE and analysed by immunoblot.

### GST pulldown

GST pulldown experiments were carried out in 96-well flat-bottomed plates (Greiner) using magnetic glutathione beads (Thermo Scientific) to capture the GST-tagged pUL21 (bait protein). First, 0.5 nmol of purified bait protein was incubated with the beads at 4°C for 30 min. The beads were washed three times with wash buffer (20 mM Tris pH 8.5, 200 mM NaCl, 0.1% NP-40, 1 mM DTT, and 1 mM EDTA), then incubated at 4°C for 60 min with various truncations of myc-CERT_L_ (prey proteins), expressed using the TNT SP6 High Yield Wheat Germ *in vitro* transcription/translation system (Promega) in accordance with the manufacturer’s protocol, followed by four washes in wash buffer. Protein was eluted using wash buffer supplemented with 50 mM reduced glutathione before being analysed by SDS-PAGE and immunoblotting.

### Antibodies

The following antibodies with listed dilutions were used for immunoblotting: rabbit anti-CERT 1:10,000 (Abcam, ab72536), mouse anti-pUL21 1:50 [1F10-D12] [18] for Fig 1F [note: this monoclonal antibody does not recognize pUL21^V382E^], rabbit anti-pUL21 1:5000 [10] for Fig 5A and S3A, mouse anti-GAPDH 1:10,000 (GeneTex, GTX28245), mouse anti-Myc 1:4000 (Millipore, 05-724), mouse anti-PP1α 1:1000 (Santa Cruz, sc-271762), rabbit anti-pUL16 1:2000 [73], mouse anti-VP5 1:50 [DM165] [74], rabbit anti-phospho-Akt substrates 1:1000 (Cell Signalling, 9611). Fluorescently labelled secondary antibodies were used at 1:10,000 dilution: LI-COR IRDye 680T donkey anti-rabbit (926–68023) and goat anti-mouse (926–68020), LI-COR IRDye 800CW conjugated donkey anti-rabbit (926–32213) and goat anti-mouse (926–32210). For immunocytochemistry, mouse anti-pUL21 1:1 [L1E4-C10], which was generated in the same immunisation experiments as anti-pUL21 [1F10-D12] [18], and Alexa Fluor 488 conjugated goat anti-mouse 1:1000 (Invitrogen, A21236) were used. For visualising HSV-1 plaques mouse anti-gD (LP2) 1:50 [75] and HRP-conjugated rabbit anti-mouse 1:5000 (DaKo, P0161) were used.

### Recombinant protein purification following bacterial expression

All recombinant proteins were expressed in *Escherichia coli* T7 Express *lysY/I^q^* cells (New England Biolabs). Except for GST-PP1γ, cells were grown in 2×TY medium at 37°C to an OD_600_ of 0.8–1.2 before cooling to 22°C and inducing protein expression by addition of 0.4 mM isopropyl β-D-thiogalactopyranoside (IPTG). At 16–20 h post-induction, cells were harvested by centrifugation and pellets were stored at -70°C until required. For GST-PP1γ, the 2×TY medium was supplemented with 1 mM MnCl_2_ and the cultures were cooled to 18°C upon reaching an OD_600_ of 0.8, followed by induction using 1 mM IPTG. For all recombinant proteins, cells were resuspended in lysis buffer (see below) at 4°C before lysis using a TS series cell disruptor (Constant Systems) at 24 kpsi. Lysates were cleared by centrifugation (40,000×g, 30 min, 4°C) and incubated with the relevant affinity resins for 1 h at 4°C before extensive washing (≥ 20 column volumes) and elution using the relevant elution buffer (see below). Samples were concentrated and subjected to size exclusion chromatography (SEC, see below). Fractions containing the desired protein as assessed by SDS-PAGE were pooled, concentrated, snap-frozen in liquid nitrogen and stored at -70°C.

The lysis buffer for pUL21-GST (20 mM Tris pH 8.5, 300 mM NaCl, 0.5 mM MgCl_2_, 1.4 mM β- mercaptoethanol, 0.05% TWEEN-20) and GST-PP1γ (50 mM Tris pH 7.5, 500 mM NaCl, 1 mM MnCl_2_, 0.5 mM MgCl_2_, 1.4 mM β-mercaptoethanol, 0.05% TWEEN-20) was supplemented with 200–400 U bovine DNase I (Merck) and 200 μL EDTA-free protease inhibitor cocktail (Merck). Cleared lysates were incubated with glutathione Sepharose 4B (Cytiva), washed with wash buffer (20 mM Tris pH 8.5 [pUL21- GST] or 7.5 [GST-PP1γ], 500 mM NaCl, 1 mM DTT, plus 1 mM MnCl_2_ [GST-PP1γ only]) and the proteins eluted using wash buffer supplemented with 25 mM reduced glutathione. SEC was performed using a HiLoad Superdex S200 16/600 column (Cytiva) equilibrated in 20 mM Tris pH 8.5, 500 mM NaCl, 1 mM DTT (pUL21-GST) or 50 mM Tris pH 7.5, 100 mM NaCl, 1 mM DTT (GST-PP1γ).

pUL21-H_6_, pUL21^V382E^-H_6_ and H_6_-pUL21C were purified in Tris buffer at pH 8.5 in with 500 mM NaCl and H_6_-miniCERT_L_ was purified in Tris buffer at pH 7.5 with 150 mM NaCl. Lysis buffer (20 mM Tris, 20 mM imidazole, NaCl, 0.5 mM MgCl_2_, 1.4 mM β-mercaptoethanol, 0.05% TWEEN-20) was supplemented with 200–400 U bovine DNase I (Merck) and 200 μL EDTA-free protease inhibitor cocktail (Merck). Cleared lysates were incubated with Ni-NTA agarose (Qiagen), washed with wash buffer (20 mM Tris, 20 mM imidazole, NaCl) and eluted using elution buffer (20 mM Tris, 250 mM imidazole, NaCl) SEC was performed using a HiLoad Superdex S200 column (Cytiva) equilibrated in 20 mM Tris, NaCl, 1 mM DTT.

### Recombinant protein purification following mammalian cell expression

StrepII-CERT^P^ used for phosphatase assays was purified following transient transfection of Freestyle 293F cells, as described before [18]. StrepII-CERT_L_^P^ and StrepII-CERT_L_^S132A^ were purified from stably transfected Freestyle 293F cells following induction with 2 μg/mL doxycycline (Fisher) for 72 hours. Next, cells were harvested by centrifugation (220×g, 5 min, 4°C) and washed once with ice-cold PBS before being resuspended in ice-cold lysis buffer with phosphatase inhibitors [StrepII-CERT_L_^P^ only] (100 mM Tris pH 8.0, 150 mM NaCl, 0.5 mM EDTA, 1 mM DTT, [10 mM tetrasodium pyrophosphate, 100 mM NaF, 17.5 mM β-glycerophosphate]). Cells were lysed by passage through a 23G needle six times and lysates were clarified by centrifugation (40,000 g, 30 min, 4°C). The supernatants were sonicated at 50% amplitude for 60 s using a sonicating probe (MSE) and the supernatants were passed through a 0.45 μm syringe filter (Sartorius). StrepII-tagged proteins were captured using 1 mL StrepTrap HP column (Cytiva) that had been pre-equilibrated in wash buffer (100 mM Tris pH 8.0, 150 mM NaCl, 0.5 mM EDTA, 1 mM DTT). After extensive washing (20 column volumes) the protein was eluted using wash buffer supplemented with 2.5 mM desthiobiotin. Pooled eluate was applied to a Superose 6 10/300 GL column (Cytiva) column equilibrated in ITC buffer (20 mM Tris pH 8.5, 500 mM NaCl, 0.5 mM Tris(2-carboxyethyl)phosphine [TCEP]) and fractions containing StrepII-CERT_L_ as assessed by SDS-PAGE were pooled, concentrated and used for downstream applications.

### Mutagenesis of viral genomes and generation of recombinant HSV-1

All HSV-1 strain KOS viruses used in this study were reconstituted from a bacterial artificial chromosome (BAC) [76] and the mutated strain was generated using the two-step Red recombination method [52] with the following primers:

Forward: 5′-CGGCTCGTAGGCCGGTACACACAGCGCCACGGCCTGTACG**AA**CCTCGGCCCGACG ACCCAGTAGGATGACGACGATAAGTAGGG

Reverse: 5′-CGTTGATGGCATCGGCCAAGACTGGGTCGTCGGGCCGAGG**TT**CGTACAGGCCGTG GCGCTGTCAACCAATTAACCAATTCTGATTAG

The generation of pUL21 deletion mutant (ΔpUL21) was described previously [18]. To generate the P0 stocks, Vero cells were transfected with the recombinant BAC DNA together with pGS403 encoding Cre recombinase (to excise the BAC cassette) using TransIt-LT1 (Mirus) following the manufacturer’s instructions. After 3 days the cells were scraped into the media, sonicated at 50% power for 30 s in a cup-horn sonicator (Branson), and titrated on Vero cell monolayers. The subsequent stocks were generated by infecting either Vero (HSV-1 WT) or HaCaT pUL21 cells (HSV-1 mutants) at MOI of 0.01 for 3 days. The cells were then scraped and isolated by centrifugation at 1,000×g for 5 min. Pellets were resuspended in 1 mL of complete DMEM supplemented with 100 U/mL penicillin, 100 μg/mL streptomycin and freeze/thawed thrice at -70°C before being aliquoted, titered on Vero cell monolayers, and stored at -70°C until required. The presence of the desired mutation in the reconstituted virus genomes was confirmed by sequencing the pUL21 gene.

### Metabolic labelling and lipid extraction for thin layer chromatography

Metabolic labelling was performed using sub-confluent (60–80% confluence) HaCaT or HaCaT21 cells grown in a 6-well plate. For analysis of stable cell lines, the cells were pre-treated for 30 min with complete DMEM containing 1 μM N-(3-Hydroxy-1-hydroxymethyl-3-phenylpropyl)dodecanamide (HPA-12, Tokyo Chemical Industry) dissolved in 0.1% dimethyl sulfoxide (DMSO, Merck), or 0.1% DMSO alone, and HPA-12 or DMSO were retained at the same concentrations throughout the subsequent incubation steps. For infection, cells were infected (below) 14 hours before metabolic labelling.

For metabolic labelling, wells were washed twice with warm PBS before incubation in 500 μL pre-warmed DMEM with 1% (v/v) Nutridoma (Merck) supplemented with 5 μM clickable sphingosine (Cayman Chemical) for 5 min (pulse). Next, cells were washed twice with warm PBS and 1 mL of pre-warmed DMEM with 1% (v/v) Nutridoma was added to each well, followed by incubation at 37°C for the indicated times of chase. Click-Sph was stored as a 3.3 mM ethanolic stock solution at -20°C.

At the indicated times of chase, the plate with cells was transferred onto the ice, washed twice with 1 mL ice-cold PBS, scraped into 300 μL of ice-cold PBS and transferred into appropriate 1.5 mL microcentrifuge tubes containing 600 μL of methanol. To each tube 150 μL of chloroform was added, followed by vigorous vortexing. The precipitated protein was pelleted (20,000×g, 1 min) at room temperature (RT). The organic supernatant was transferred to separate 2 mL tubes containing 300 μL chloroform. 600 μL of 0.1% acetic acid in H_2_O was subsequently added to each tube to induce formation of two phases. Following extensive vortexing, the phases were separated by centrifugation (20,000×g, 1 min, RT) and the lower phase was transferred to a new 1.5 mL microcentrifuge tube. This lipid-containing solvent phase was dried in a UniVapo centrifugal vacuum concentrator (UniEquip) at 30°C for 20 min.

If required for SDS-PAGE analysis, the protein pellet was solubilized with 5 μL 10% (w/v) SDS, diluted in 75 μL lysis buffer (20 mM Tris 7.5, 150 mM NaCl, 1% (v/v) Triton X-100), boiled for 10 min and sonicated at 75% power for 60 s in a cup-horn sonicator (Branson). The protein samples were mixed with 5× loading buffer before being subjected to SDS-PAGE analysis.

### Click reaction and thin layer chromatography of labelled lipids

Lipid standards (16:0(Alkyne)-18:1 PC [Avanti], pacFA GalCer [Avanti], d18:1(Alkyne)-6:0 Cer [Cayman Chemicals], 18:0(Alkyne) Sph [Cayman Chemicals]) or extracted lipids (above) were resuspended in 20 μL of 1:1 chloroform:ethanol. To each tube 1 μL of Tris((1-benzyl-4-triazolyl)methyl)amine (TBTA) solution (2.5 mM in DMSO; Merck), 10 μL of Tetrakis(acetonitrile)copper(I) tetrafluoroborate (10 mM in acetonitrile; Merck) and 1 μL of 3-azido-7-hydroxycoumarin solution (1 mM in EtOH; Jena Bioscience) was added. After brief mixing, the reaction was dried in a UniVapo centrifugal vacuum concentrator (UniEquip) for 10 min at 45°C. Lipids were dissolved in 20 μL of 65:25:4:1 chloroform:methanol:water:acetic acid, of which 8 μL was spotted on a 20×10 cm HPTLC Silica gel 60 plates (Merck). When dried, the plate was developed for 5 cm with 65:25:4:1 chloroform:methanol:water:acetic acid, dried again and developed for 9 cm with 1:1 hexane:ethylacetate. Labelled lipids were visualized via UV illumination using a G:BOX Chemi XX9 (Syngene), bands were quantified using the Image Studio Lite software (LI-COR) with local background subtraction, and statistical tests were performed using Prism 7 (GraphPad Software).

### Virus infections

Monolayers of indicated cells were infected by overlaying with the appropriate viruses, diluted in complete DMEM to the specified MOI. The time of addition was designated 0 hours post-infection (hpi). Cells were incubated with the inoculum at 37°C in a humidified 5% CO_2_ atmosphere for 1 hour, rocking the tissue culture plate housing the monolayers every 15 min, before complete DMEM supplemented with 100 U/mL penicillin, 100 μg/mL streptomycin was added to dilute the original infection medium five- fold. Infected cells were incubated at 37°C in a humidified 5% CO_2_ atmosphere and harvested at the specified times.

### Immunoblotting

For analysis of protein expression and phosphorylation, cells were washed two times with ice-cold 50 mM Tris pH 8.5, 150 mM NaCl and scraped into 100 μL of ice-cold lysis buffer (50 mM Tris pH 8.5, 150mM NaCl, 1% (v/v) Triton-X100, 1% (v/v) EDTA-free protease inhibitor cocktail [Merck], 10 mM tetrasodium pyrophosphate, 100 mM NaF and 17.5 mM β-glycerophosphate). After 30 min incubation on ice, the lysates were sonicated (two 15 s pulses at 50% power) in a cup-horn sonicator (Branson) followed by centrifugation at 20,000×g for 10 min. For all samples, protein concentrations were determined using a BCA assay (Peirce) and normalized protein lysates were analysed by SDS-PAGE. For enhanced separation of phosphorylated proteins, 7% (w/v) acrylamide gels contained 50 μM MnCl_2_ and 25 μM PhosTag reagent (Wako) where indicated. Separated proteins were then transferred to nitrocellulose membranes using the Mini-PROTEAN system (Bio-Rad) and analysed by immunoblotting, signals being detected using Odyssey CLx Imaging System (LI-COR). For quantitation of CERT phosphoform relative abundance, the signal for CERT^O^ band alone and for all CERT bands (CERT^O^ + CERT^P^) were quantitated using Image Studio Lite software (LI-COR) with local background subtraction. Statistical tests were performed using Prism 7 (GraphPad Software).

### Analytical SEC of miniCERT_L_:pUL21C complex

Analytical SEC experiments were performed using Superdex 200 10/300 GL (Cytiva) column equilibrated in 20 mM Tris pH 8.5, 500 mM NaCl, 1 mM DTT. To allow complex formation, 200 µL of 140 µM H_6_-miniCERT_L_ and 680 µL of 55.4 µM H_6_-pUL21C were mixed and incubated for 15 min at room temperature prior to injection.

### Isothermal titration calorimetry (ITC)

ITC experiments were performed at 25°C using an MicroCal PEAQ-ITC automated calorimeter (Malvern Panalytical). Proteins were transferred into ITC buffer (20 mM Tris pH 8.5, 500 mM NaCl, 0.5 mM TCEP) either by SEC or extensive dialysis prior to experiments. Titrants (StrepII-CERT_L_^P^ at 192–328 μM, StrepII-CERT_L_^S132A^ at 196–334 μM or H_6_-miniCERT_L_ at 195–376 μM) were titrated into titrates (pUL21- H_6_ at 20–33 μM, pUL21^V382E^-H_6_ at 19.5–32 μM or H_6_-pUL21C at 30–32.5 μM) using either 19×2 μL or 12×3 μL injections. Data were analysed using the MicroCal PEAQ-ITC analysis software (Malvern Panalytical) and fitted using a one-site binding model.

### Multi-angle light scattering (MALS)

MALS data were collected immediately following SEC (SEC-MALS) by inline measurement of static light scattering (DAWN 8+; Wyatt Technology), differential refractive index (Optilab T-rEX; Wyatt Technology), and UV absorbance (1260 UV; Agilent Technologies). Samples (100 μL) were injected onto an Superose 6 increase 10/300 GL column (Cytiva) equilibrated in in 20 mM Tris pH 8.5, 500 mM NaCl, 0.5 mM TCEP at 0.4 mL/min. The protein samples were injected at the concentration of 2 mg/mL. Molecular masses were calculated using ASTRA 6 (Wyatt Technology) using a protein dn/dc of 0.186 mL/g.

### Small-angle X-ray scattering (SAXS)

SAXS experiments were performed in batch mode (H_6_-pUL21C and H_6_-miniCERT_L_) or inline following SEC (H_6_-pUL21C:H_6_-miniCERT_L_) at EMBL-P12 bioSAXS beam line (PETRAIII, DESY, Hamburg, Germany) [77, 78].

For SAXS in batch mode, scattering data (*I*(*s*) versus *s*, where *s* = 4πsinθ/λ nm^-1^, 2θ is the scattering angle, and λ is the X-ray wavelength, 0.124 nm) were collected from the indicated batch samples and a corresponding solvent blank (20 mM Tris pH 8.5, 500 mM NaCl, 3% (v/v) glycerol, 1 mM DTT). The scattering profiles of H_6_-miniCERT_L_ (1.73, 2.59, 3.46 and 6.91 mg/mL) and H_6_-pUL21C (0.76 mg/mL) were measured in continuous-flow mode using an automated sample changer (30 μL sample at 20°C; 1 mm pathlength). The sample and buffer measurements were measured as 100 ms data frames on a Pilatus 6M area detector for total exposure times of 2.1 and 2.6 s, respectively. For H_6_-miniCERT_L_, the resulting SAXS curves were extrapolated to infinite dilution.

For SEC-SAXS experiments, the scattering data were recorded using a Pilatus 6M detector (Dectris) with 1 s sample exposure times for a total of 3600 data frames spanning the entire course of the SEC separation. Proteins were mixed at 1:1 molar ratio, incubated on ice for 15 min, and then concentrated using a centrifugal concentrator to the desired concentration as estimated from absorbance using extinction coefficients [79] calculated assuming a 1:1 molar ratio. 40 μL of purified H_6_-pUL21C:H_6_- miniCERT_L_ (4.3 mg/mL) was injected at 0.35 mL/min onto an Superdex 200 Increase 5/150 GL column (Cytiva) equilibrated in 20 mM Tris pH 8.5, 500 mM NaCl, 3% (v/v) glycerol, 1 mM DTT. SAXS data were recorded from a single peak (frames 1245–1296 s) and solvent blank was collected from pre-elution frames (frames 247–411). Primary data reduction was carried out using CHROMIXS [80] and 2D-to-1D radial averaging was performed using the SASFLOW pipeline [81]. Processing and analysis of the SAXS output was performed using the ATSAS 3.0.2 software package [82]. The extrapolated forward scattering intensity at zero angle, *I*(0), and the radius of gyration, *R_g_*, were calculated from the Guinier approximation (ln*I*(*s*) vs *s*^2^, for *sR_g_* < 1.3). The maximum particle dimension, D_max_, was estimated based on the probable distribution of real-space distances *p*(*r*) which was calculated using GNOM [83]. A concentration-independent estimate of molecular weight was determined using a Bayesian consensus method [84]. All structural parameters are reported in Table S1.

*Ab initio* modelling was performed using GASBOR [49] and figures show the models that best fit their corresponding SAXS profiles (lowest χ^2^). Pseudo-atomic modelling was performed using CORAL [85]. For H_6_-miniCERT_L_, crystal structures of the CERT PH (PDB ID 4HHV) [24] and START (PDB ID 2E3M) [25] domains were used, the latter including predicted secondary structure as modelled by I-TASSER [86] for intron 11 (residues 371-396), present only in CERT_L_. The miniCERT_L_ 23 amino acid linker and the purification tag (MAH_6_) were modelled using CORAL. The model with the best fit was superimposed onto the PH:START crystal structure (PDB ID 5JJD) [46] by aligning the START domains using PyMOL version 2.5.2 (Schrödinger). For H_6_-pUL21C, the N-terminal (MAH_6_QD) and C-terminal (HGQSV) elements missing from the crystal structure (PDB ID 5ED7) [50] were modelled using CORAL. For H_6_- pUL21C:H_6_-miniCERT_L_, the heterodimer was modelled using the SAXS curve of the complex using a fixed conformation of H_6_-pUL21C and mobile CERT domains, with unstructured elements modelled as above. All the models were assessed for fit to the corresponding SAXS profiles (χ^2^ and CorMap) using CRYSOL [87].

### Differential scanning fluorimetry

Differential scanning fluorimetry experiments were carried out using Viia7 real-time PCR system (Applied Biosystems) and 1× Protein Thermal Shift dye (Applied Biosystems). The assay buffer (20 mM Tris pH 8.5, 500 mM NaCl, 1 mM DTT) was mixed with dye stock solution and protein solution (H_6_- pUL21 or H_6_-pUL21^V382E^ in an 8:1:1 ratio, giving 1 ng protein in a final volume of 20 μL, and measurements were performed in triplicate. Samples were heated from 25 to 95°C at 1 degree per 20 s and fluorescence was monitored at each increment. The melting temperature (T_m_) is the inflection point of the sigmoidal melting curve, determined by non-linear curve fitting to the Boltzmann equation using Prism 7 (GraphPad Software).

### In vitro dephosphorylation assays

Dephosphorylation assays were performed upon 0.5 μM of purified CERT^P^ using a fixed concentration of GST-PP1γ (25 nM) in the presence of pUL21-H_6_ or pUL21^V382E^ -H_6_ at the indicated concentrations. Reactions (50 μL) proceeded in assay buffer (150 mM NaCl, 20 mM Tris pH 8.5, 0.1% TWEEN-20, 1 mM MnCl_2_) for 30 min at 30°C before being stopped by the addition of 50 μL of 2× SDS-PAGE loading buffer and boiling at 95°C for 5 min. Samples were analysed by SDS-PAGE using 7% (w/v) acrylamide gels supplemented with 25 μM PhosTag reagent and 50 μM MnCl_2_ and the protein was visualized using InstantBlue Coomassie stain (Expedeon). To measure the ratio of CERT^O^ to total CERT, Coomassie-stained gels were scanned using an Odyssey CLx Imaging System (LI-COR). The signals detected in the 700 nm channel for the CERT^O^ band alone and for all CERT bands (CERT^O^ + CERT^P^) were quantitated using Image Studio Lite software (LI-COR) with local background subtraction.

### Plaque size analysis

Confluent monolayers of Vero or HaCaT cells in 6-well tissue culture plates were infected with 100 plaque forming units of the indicated virus diluted in complete DMEM to a final volume of 500 μL. Following the adsorption for 1 h at 37°C in a humidified 5% CO_2_ atmosphere, rocking the plate every 15 min, the cells were overlaid with plaque assay media (DMEM supplemented with 0.3% high viscosity carboxymethyl cellulose, 0.3% low viscosity carboxymethyl cellulose, 2% (v/v) FCS, 2 mM L-glutamine, 100 U/mL penicillin and 100 μg/mL streptomycin) and incubated for further 48 hours. Next, cells were fixed with 3.7% (v/v) formal saline for 20 min, washed three times with PBS and incubated for 1 h with mouse anti-gD (LP2) [75], diluted 1:50 in blocking buffer (PBS supplemented with 1% (w/v) BSA and 0.1% TWEEN-20). Cells were washed thrice in PBS and incubated for 30 min with HRP-conjugated rabbit anti-mouse antibody (DaKo, P0161) diluted 1:5000 in blocking buffer, followed by two subsequent PBS washes and one wash with ultrapure water. Plaques were visualized using the TrueBlue peroxidase substrate in accordance with the manufacturer’s instructions (Seracare). Plaques were scanned at 1200 dpi and plaque areas were measured using Fiji [88, 89].

### Single-step (high MOI) virus growth assays

Virus growth assays were performed in technical duplicate for each independent experiment. Monolayers of HaCaT or Vero cells were infected with the indicated viruses diluted in complete DMEM to an MOI of 10. The time of virus addition was designated 0 hpi. After adsorption for 1 h at 37°C in a humidified 5% CO_2_ atmosphere, rocking the plates every 15 min, extracellular virus particles were neutralized with an acid wash (40 mM citric acid, 135 mM NaCl, 10 mM KCl, pH 3.0) for 1 min and cells were then washed three times with PBS before being subsequently overlaid with complete DMEM. At designated times post-infection, cells were harvested by freezing the plate at -70°C. When all plates were frozen, samples were freeze/thawed one more time before scraping and transferring to 1.5 mL tubes and storing at -70°C until titration. Titrations were performed on monolayers of Vero cells. Serial dilutions of the samples were used to inoculate the cells for 1 h, followed by overlaying with DMEM containing 0.3% high viscosity carboxymethyl cellulose, 0.3% low viscosity carboxymethyl cellulose, 2% (v/v) FBS, 2 mM L-glutamine, 100 U/mL penicillin and 100 μg/mL streptomycin. Three days later, cells were fixed in 3.7% (v/v) formal saline for 20 min, washed with water and stained with 0.1% toluidine blue. Statistical tests were performed using Prism 7 (GraphPad Software).

### Virus release assays

Monolayers of HaCaT or Vero cells were infected as described above for single-step virus growth assays, infections being performed in technical duplicate or triplicate for each independent experiment. At 12 hpi, the media (500 μL) were harvested to 1.5 mL Eppendorf tubes, spun down for 5 min at 1000 g to remove any detached cells, 300 μL of the supernatant was carefully transferred to fresh tubes and stored at -70°C until titration. The cells were overlaid with 500 μL of fresh DMEM and the plates were immediately frozen at -70°C. Titrations were performed on monolayers of Vero cells as described above. Statistical tests were performed using Prism 7 (GraphPad Software).

### Immunocytochemistry

Cells grown on #1.5 coverslips were infected at an MOI of 1 as described above. At 14 hpi, cells were washed with PBS and incubated with freezing cold (-20°C) methanol for 5 min at -20°C. Coverslips were washed with PBS, followed by incubation with blocking buffer (1% (w/v) BSA in PBS) for 30 min at RT. Primary antibodies (above) were diluted in blocking buffer and incubated with coverslips for 1 h. Coverslips were washed ten times with PBS before incubation for 45 min with the secondary antibodies (above) diluted in blocking buffer. Coverslips were washed ten times in PBS and ten times in ultrapure water before mounting on slides using Mowiol 4–88 (Merck) containing 200 nM 4′,6-diamidino-2- phenylindole (DAPI). Images were acquired using an inverted Olympus IX81 widefield microscope with a 60× Plan Apochromat N oil objective (numerical aperture 1.42) (Olympus) and Retiga EXi Fast1394 interline CCD camera (QImaging).

## Acknowledgements

The synchrotron SAXS data was collected at beamline P12 operated by EMBL Hamburg at the PETRA III storage ring (DESY, Hamburg, Germany). We thank Chris Boutell (MRC-University of Glasgow Centre for Virus Research), John Willis (University of Pennsylvania) and David Ron (Cambridge Institute for Medical Research) for kindly providing antibodies or plasmids. This work was funded by a Royal Society University Research Fellowship (UF100371), a Royal Society Enhancement Award (RGF\EA\180151) and a Wellcome Trust Senior Research Fellowship (219447/Z/19/Z) to JED, a Biotechnology and Biological Sciences Research Council (BBSRC) Research Grant (BB/M021424/1) and a Medical Research Council Research Grant (MR/T016493/1) to CMC, and by a Sir Henry Dale Fellowship, jointly funded by the Wellcome Trust and the Royal Society (098406/Z/12/B) to SCG. DIS and CMJ acknowledge the support of iNEXT-Discovery (project number 871037) funded by the Horizon 2020 programme of the European Commission. The funders had no role in study design, data collection and analysis, decision to publish, or preparation of the manuscript. For the purpose of open access, the authors have applied a Creative Commons Attribution (CC BY) licence to any Author Accepted Manuscript version arising from this submission.

## Data availability

The SAXS data measured for each concentration, with an accompanying report, are available in the Small Angle Scattering Biological Data Bank (SASBDB) [90], entries SASDNB7, SASDNC7 and SASDND7.

## References

1. Looker KJ, Magaret AS, May MT, Turner KME, Vickerman P, Gottlieb SL, et al. Global and Regional Estimates of Prevalent and Incident Herpes Simplex Virus Type 1 Infections in 2012. PloS One. 2015;10: e0140765. doi:10.1371/journal.pone.0140765

2. Kulej K, Avgousti DC, Sidoli S, Herrmann C, Fera AND, Kim ET, et al. Time-resolved Global and Chromatin Proteomics during Herpes Simplex Virus Type 1 (HSV-1) Infection. Mol Cell Proteomics MCP. 2017;16: S92. doi:10.1074/mcp.M116.065987

3. Soh TK, Davies CTR, Muenzner J, Hunter LM, Barrow HG, Connor V, et al. Temporal Proteomic Analysis of Herpes Simplex Virus 1 Infection Reveals Cell-Surface Remodeling via pUL56- Mediated GOPC Degradation. Cell Rep. 2020;33: 108235. doi:10.1016/j.celrep.2020.108235

4. Scherer KM, Manton JD, Soh TK, Mascheroni L, Connor V, Crump CM, et al. A fluorescent reporter system enables spatiotemporal analysis of host cell modification during herpes simplex virus-1 replication. J Biol Chem. 2021;296: 100236. doi:10.1074/jbc.RA120.016571

5. Nahas KL, Connor V, Scherer KM, Kaminski CF, Harkiolaki M, Crump CM, et al. Near-native state imaging by cryo-soft-X-ray tomography reveals remodelling of multiple cellular organelles during HSV-1 infection. PLoS Pathog. 2022;18: e1010629. doi:10.1371/journal.ppat.1010629

6. Bigalke JM, Heldwein EE. Nuclear Exodus: Herpesviruses Lead the Way. Annu Rev Virol. 2016;3: 387–409. doi:10.1146/annurev-virology-110615-042215

7. Owen DJ, Crump CM, Graham SC. Tegument Assembly and Secondary Envelopment of Alphaherpesviruses. Viruses. 2015;7: 5084–5114. doi:10.3390/v7092861

8. Cocchi F, Menotti L, Dubreuil P, Lopez M, Campadelli-Fiume G. Cell-to-Cell Spread of Wild-Type Herpes Simplex Virus Type 1, but Not of Syncytial Strains, Is Mediated by the Immunoglobulin-Like Receptors That Mediate Virion Entry, Nectin1 (PRR1/HveC/HIgR) and Nectin2 (PRR2/HveB). J Virol. 2000;74: 3909–3917. doi:10.1128/JVI.74.8.3909-3917.2000

9. Le Sage V, Jung M, Alter JD, Wills EG, Johnston SM, Kawaguchi Y, et al. The herpes simplex virus 2 UL21 protein is essential for virus propagation. J Virol. 2013;87: 5904–5915. doi:10.1128/JVI.03489-12

10. Harper AL, Meckes DG, Marsh JA, Ward MD, Yeh P-C, Baird NL, et al. Interaction domains of the UL16 and UL21 tegument proteins of herpes simplex virus. J Virol. 2010;84: 2963–2971. doi:10.1128/JVI.02015-09

11. Klupp BG, Böttcher S, Granzow H, Kopp M, Mettenleiter TC. Complex formation between the UL16 and UL21 tegument proteins of pseudorabies virus. J Virol. 2005;79: 1510–1522. doi:10.1128/JVI.79.3.1510-1522.2005

12. Han J, Chadha P, Starkey JL, Wills JW. Function of glycoprotein E of herpes simplex virus requires coordinated assembly of three tegument proteins on its cytoplasmic tail. Proc Natl Acad Sci. 2012;109: 19798–19803. doi:10.1073/pnas.1212900109

13. Takakuwa H, Goshima F, Koshizuka T, Murata T, Daikoku T, Nishiyama Y. Herpes simplex virus encodes a virion-associated protein which promotes long cellular processes in over-expressing cells. Genes Cells. 2001;6: 955–966. doi:10.1046/j.1365-2443.2001.00475.x

14. Yan K, Liu J, Guan X, Yin Y-X, Peng H, Chen H-C, et al. The Carboxyl Terminus of Tegument Protein pUL21 Contributes to Pseudorabies Virus Neuroinvasion. J Virol. 2019;93: 02052–18. doi:10.1128/JVI.02052-18

15. Finnen RL, Banfield BW. CRISPR/Cas9 Mutagenesis of UL21 in Multiple Strains of Herpes Simplex Virus Reveals Differential Requirements for pUL21 in Viral Replication. Viruses. 2018;10. doi:10.3390/v10050258

16. Sarfo A, Starkey J, Mellinger E, Zhang D, Chadha P, Carmichael J, et al. The UL21 Tegument Protein of Herpes Simplex Virus 1 Is Differentially Required for the Syncytial Phenotype. J Virol. 2017;91: 01161–17. doi:10.1128/JVI.01161-17

17. Klupp BG, Lomniczi B, Visser N, Fuchs W, Mettenleiter TC. Mutations affecting the UL21 gene contribute to avirulence of pseudorabies virus vaccine strain Bartha. Virology. 1995;212: 466–473. doi:10.1006/viro.1995.1504

18. Benedyk TH, Muenzner J, Connor V, Han Y, Brown K, Wijesinghe KJ, et al. pUL21 is a viral phosphatase adaptor that promotes herpes simplex virus replication and spread. PLOS Pathog. 2021;17: e1009824. doi:10.1371/journal.ppat.1009824

19. Peti W, Nairn AC, Page R. Structural basis for protein phosphatase 1 regulation and specificity. FEBS J. 2013;280: 596–611. doi:10.1111/j.1742-4658.2012.08509.x

20. Muradov JH, Finnen RL, Gulak MA, Hay TJM, Banfield BW. pUL21 regulation of pUs3 kinase activity influences the nature of nuclear envelope deformation by the HSV-2 nuclear egress complex. PLoS Pathog. 2021;17: e1009679. doi:10.1371/journal.ppat.1009679

21. Fukasawa M, Nishijima M, Hanada K. Genetic evidence for ATP-dependent endoplasmic reticulum-to-Golgi apparatus trafficking of ceramide for sphingomyelin synthesis in Chinese hamster ovary cells. J Cell Biol. 1999;144: 673–685. doi:10.1083/jcb.144.4.673

22. Funakoshi T, Yasuda S, Fukasawa M, Nishijima M, Hanada K. Reconstitution of ATP- and cytosol- dependent transport of de novo synthesized ceramide to the site of sphingomyelin synthesis in semi-intact cells. J Biol Chem. 2000;275: 29938–29945. doi:10.1074/jbc.M004470200

23. Hanada K, Kumagai K, Yasuda S, Miura Y, Kawano M, Fukasawa M, et al. Molecular machinery for non-vesicular trafficking of ceramide. Nature. 2003;426: 803–809. doi:10.1038/nature02188

24. Prashek J, Truong T, Yao X. Crystal Structure of the Pleckstrin Homology Domain from the Ceramide Transfer Protein: Implications for Conformational Change upon Ligand Binding. PLoS ONE. 2013;8: e79590. doi:10.1371/journal.pone.0079590

25. Kudo N, Kumagai K, Tomishige N, Yamaji T, Wakatsuki S, Nishijima M, et al. Structural basis for specific lipid recognition by CERT responsible for nonvesicular trafficking of ceramide. Proc Natl Acad Sci. 2008;105: 488–493. doi:10.1073/pnas.0709191105

26. Charruyer A, Bell SM, Kawano M, Douangpanya S, Yen T-Y, Macher BA, et al. Decreased Ceramide Transport Protein (CERT) Function Alters Sphingomyelin Production following UVB Irradiation. J Biol Chem. 2008;283: 16682–16692. doi:10.1074/jbc.M800799200

27. Kawano M, Kumagai K, Nishijima M, Hanada K. Efficient trafficking of ceramide from the endoplasmic reticulum to the Golgi apparatus requires a VAMP-associated protein-interacting FFAT motif of CERT. J Biol Chem. 2006;281: 30279–30288. doi:10.1074/jbc.M605032200

28. Kumagai K, Kawano M, Shinkai-Ouchi F, Nishijima M, Hanada K. Interorganelle trafficking of ceramide is regulated by phosphorylation-dependent cooperativity between the PH and START domains of CERT. J Biol Chem. 2007;282: 17758–17766. doi:10.1074/jbc.M702291200

29. Hannun YA, Obeid LM. Sphingolipids and their metabolism in physiology and disease. Nat Rev Mol Cell Biol. 2018;19: 175–191. doi:10.1038/nrm.2017.107

30. Sezgin E, Levental I, Mayor S, Eggeling C. The mystery of membrane organization: composition, regulation and roles of lipid rafts. Nat Rev Mol Cell Biol. 2017;18: 361–374. doi:10.1038/nrm.2017.16

31. Olsen ASB, Faergeman NJ. Sphingolipids: membrane microdomains in brain development, function and neurological diseases. Open Biol. 2017;7: 170069. doi:10.1098/rsob.170069

32. Elwell CA, Engel JN. Lipid acquisition by intracellular Chlamydiae. Cell Microbiol. 2012;14: 1010– 1018. doi:10.1111/j.1462-5822.2012.01794.x

33. Gewaid H, Aoyagi H, Arita M, Watashi K, Suzuki R, Sakai S, et al. Sphingomyelin Is Essential for the Structure and Function of the Double-Membrane Vesicles in Hepatitis C Virus RNA Replication Factories. J Virol. 2020;94: e01080–20. doi:10.1128/JVI.01080-20

34. Ray EK, Blough HA. The effect of herpesvirus infection and 2-deoxy-d-glucose on glycosphingolipids in BHK-21 cells. Virology. 1978;88: 118–127. doi:10.1016/0042-6822(78)90115-0

35. Steinhart WL, Busch JS, Oettgen JP, Howland JL. Sphingolipid metabolism during infection of human fibroblasts by herpes simplex virus type 1. Intervirology. 1984;21: 70–76. doi:10.1159/000149504

36. Pastenkos G, Miller JL, Pritchard SM, Nicola AV. Role of Sphingomyelin in Alphaherpesvirus Entry. J Virol. 2019;93: e01547–18. doi:10.1128/JVI.01547-18

37. Lang J, Bohn P, Bhat H, Jastrow H, Walkenfort B, Cansiz F, et al. Acid ceramidase of macrophages traps herpes simplex virus in multivesicular bodies and protects from severe disease. Nat Commun. 2020;11: 1338. doi:10.1038/s41467-020-15072-8

38. Roussel E, Lippe R. Cellular Protein Kinase D Modulators Play a Role during Multiple Steps of Herpes Simplex Virus 1 Egress. J Virol. 2018;92: e01486–18. doi:10.1128/JVI.01486-18

39. Shevchenko A, Simons K. Lipidomics: coming to grips with lipid diversity. Nat Rev Mol Cell Biol. 2010;11: 593–598. doi:10.1038/nrm2934

40. Sunshine H, Iruela-Arispe ML. Membrane Lipids and Cell Signaling. Curr Opin Lipidol. 2017;28: 408–413. doi:10.1097/MOL.0000000000000443

41. Gaebler A, Milan R, Straub L, Hoelper D, Kuerschner L, Thiele C. Alkyne lipids as substrates for click chemistry-based in vitro enzymatic assays. J Lipid Res. 2013;54: 2282–2290. doi:10.1194/jlr.D038653

42. Gerl MJ, Bittl V, Kirchner S, Sachsenheimer T, Brunner HL, Lüchtenborg C, et al. Sphingosine-1- Phosphate Lyase Deficient Cells as a Tool to Study Protein Lipid Interactions. PLOS ONE. 2016;11: e0153009. doi:10.1371/journal.pone.0153009

43. Morash SC, Cook HW, Spence MW. Lysophosphatidylcholine as an intermediate in phosphatidylcholine metabolism and glycerophosphocholine synthesis in cultured cells: an evaluation of the roles of 1-acyl- and 2-acyl-lysophosphatidylcholine. Biochim Biophys Acta. 1989;1004: 221–229. doi:10.1016/0005-2760(89)90271-3

44. Yasuda S, Kitagawa H, Ueno M, Ishitani H, Fukasawa M, Nishijima M, et al. A Novel Inhibitor of Ceramide Trafficking from the Endoplasmic Reticulum to the Site of Sphingomyelin Synthesis. J Biol Chem. 2001;276: 43994–44002. doi:10.1074/jbc.M104884200

45. Prischi F, Filippakopoulos P. Editorial: Structural Studies of Protein Complexes in Signaling Pathways. Front Mol Biosci. 2021;8: 200. doi:10.3389/fmolb.2021.641932

46. Prashek J, Bouyain S, Fu M, Li Y, Berkes D, Yao X. Interaction between the PH and START domains of ceramide transfer protein competes with phosphatidylinositol 4-phosphate binding by the PH domain. J Biol Chem. 2017;292: 14217–14228. doi:10.1074/jbc.M117.780007

47. Sugiki T, Takeuchi K, Yamaji T, Takano T, Tokunaga Y, Kumagai K, et al. Structural Basis for the Golgi Association by the Pleckstrin Homology Domain of the Ceramide Trafficking Protein (CERT). J Biol Chem. 2012;287: 33706–33718. doi:10.1074/jbc.M112.367730

48. Raya A, Revert-Ros F, Martinez-Martinez P, Navarro S, Roselló E, Vieites B, et al. Goodpasture Antigen-binding Protein, the Kinase That Phosphorylates the Goodpasture Antigen, Is an Alternatively Spliced Variant Implicated in Autoimmune Pathogenesis. J Biol Chem. 2000;275: 40392–40399. doi:10.1074/jbc.M002769200

49. Svergun DI, Petoukhov MV, Koch MHJ. Determination of Domain Structure of Proteins from X- Ray Solution Scattering. Biophys J. 2001;80: 2946–2953. doi:10.1016/S0006-3495(01)76260-1

50. Metrick CM, Heldwein EE. Novel Structure and Unexpected RNA-Binding Ability of the C-Terminal Domain of Herpes Simplex Virus 1 Tegument Protein UL21. J Virol. 2016;90: 5759–5769. doi:10.1128/JVI.00475-16

51. Metrick CM, Chadha P, Heldwein EE. The unusual fold of herpes simplex virus 1 UL21, a multifunctional tegument protein. J Virol. 2015;89: 2979–2984. doi:10.1128/JVI.03516-14

52. Tischer BK, Smith GA, Osterrieder N. En passant mutagenesis: a two step markerless red recombination system. Methods Mol Biol Clifton NJ. 2010;634: 421–430. doi:10.1007/978-1-60761-652-8_30

53. Chuluunbaatar U, Roller R, Feldman ME, Brown S, Shokat KM, Mohr I. Constitutive mTORC1 activation by a herpesvirus Akt surrogate stimulates mRNA translation and viral replication. Genes Dev. 2010;24: 2627–2639. doi:10.1101/gad.1978310

54. Roussel É, Lippé R. Cellular Protein Kinase D Modulators Play a Role during Multiple Steps of Herpes Simplex Virus 1 Egress. J Virol. 2018;92: e01486–18. doi:10.1128/JVI.01486-18

55. Bell C, Desjardins M, Thibault P, Radtke K. Proteomics analysis of herpes simplex virus type 1- infected cells reveals dynamic changes of viral protein expression, ubiquitylation, and phosphorylation. J Proteome Res. 2013;12: 1820–1829. doi:10.1021/pr301157j

56. Langeland N, Haarr L, Holmsen H. Polyphosphoinositide metabolism in baby-hamster kidney cells infected with herpes simplex virus type 1. Biochem J. 1986;237: 707–712. doi:10.1042/bj2370707

57. Sutter E, de Oliveira AP, Tobler K, Schraner EM, Sonda S, Kaech A, et al. Herpes simplex virus 1 induces de novo phospholipid synthesis. Virology. 2012;429: 124–135. doi:10.1016/j.virol.2012.04.004

58. Han X. Lipidomics for studying metabolism. Nat Rev Endocrinol. 2016;12: 668–679. doi:10.1038/nrendo.2016.98

59. Xu T, Hu C, Xuan Q, Xu G. Recent advances in analytical strategies for mass spectrometry-based lipidomics. Anal Chim Acta. 2020;1137: 156–169. doi:10.1016/j.aca.2020.09.060

60. Campadelli G, Brandimarti R, Lazzaro CD, Ward PL, Roizman B, Torrisi MR. Fragmentation and dispersal of Golgi proteins and redistribution of glycoproteins and glycolipids processed through the Golgi apparatus after infection with herpes simplex virus 1. Proc Natl Acad Sci. 1993;90: 2798–2802. doi:10.1073/pnas.90.7.2798

61. Kumagai K, Hanada K. Structure, functions and regulation of CERT, a lipid-transfer protein for the delivery of ceramide at the ER-Golgi membrane contact sites. FEBS Lett. 2019;593: 2366–2377. doi:10.1002/1873-3468.13511

62. Rémillard-Labrosse G, Mihai C, Duron J, Guay G, Lippé R. Protein kinase D-dependent trafficking of the large Herpes simplex virus type 1 capsids from the TGN to plasma membrane. Traffic Cph Den. 2009;10: 1074–1083. doi:10.1111/j.1600-0854.2009.00939.x

63. Alonso A, Goñi FM. The Physical Properties of Ceramides in Membranes. Annu Rev Biophys. 2018;47: 633–654. doi:10.1146/annurev-biophys-070317-033309

64. Bigalke JM, Heldwein EE. Structural basis of membrane budding by the nuclear egress complex of herpesviruses. EMBO J. 2015;34: 2921–2936. doi:10.15252/embj.201592359

65. Thorsen MK, Lai A, Lee MW, Hoogerheide DP, Wong GCL, Freed JH, et al. Highly Basic Clusters in the Herpes Simplex Virus 1 Nuclear Egress Complex Drive Membrane Budding by Inducing Lipid Ordering. mBio. 12: e01548–21. doi:10.1128/mBio.01548-21

66. Ogretmen B. Sphingolipid metabolism in cancer signalling and therapy. Nat Rev Cancer. 2018;18: 33–50. doi:10.1038/nrc.2017.96

67. Galvan V, Roizman B. Herpes simplex virus 1 induces and blocks apoptosis at multiple steps during infection and protects cells from exogenous inducers in a cell-type-dependent manner. Proc Natl Acad Sci U S A. 1998;95: 3931–3936. doi:10.1073/pnas.95.7.3931

68. Li Z, Michael IP, Zhou D, Nagy A, Rini JM. Simple piggyBac transposon-based mammalian cell expression system for inducible protein production. Proc Natl Acad Sci U S A. 2013;110: 5004– 5009. doi:10.1073/pnas.1218620110

69. Zufferey R, Donello JE, Trono D, Hope TJ. Woodchuck hepatitis virus posttranscriptional regulatory element enhances expression of transgenes delivered by retroviral vectors. J Virol. 1999;73: 2886–2892. doi:10.1128/JVI.73.4.2886-2892.1999

70. Neidel S, Maluquer de Motes C, Mansur DS, Strnadova P, Smith GL, Graham SC. Vaccinia virus protein A49 is an unexpected member of the B-cell Lymphoma (Bcl)-2 protein family. J Biol Chem. 2015;290: 5991–6002. doi:10.1074/jbc.M114.624650

71. Chen R, Rato C, Yan Y, Crespillo-Casado A, Clarke HJ, Harding HP, et al. G-actin provides substrate-specificity to eukaryotic initiation factor 2α holophosphatases. eLife. 2015;4. doi:10.7554/eLife.04871

72. Boukamp P, Petrussevska RT, Breitkreutz D, Hornung J, Markham A, Fusenig NE. Normal keratinization in a spontaneously immortalized aneuploid human keratinocyte cell line. J Cell Biol. 1988;106: 761–771. doi:10.1083/jcb.106.3.761

73. Carmichael JC, Wills JW. Differential Requirements for gE, gI, and UL16 among Herpes Simplex Virus 1 Syncytial Variants Suggest Unique Modes of Dysregulating the Mechanism of Cell-to-Cell Spread. J Virol. 2019;93. doi:10.1128/JVI.00494-19

74. McClelland DA, Aitken JD, Bhella D, McNab D, Mitchell J, Kelly SM, et al. pH reduction as a trigger for dissociation of herpes simplex virus type 1 scaffolds. J Virol. 2002;76: 7407–7417. doi:10.1128/jvi.76.15.7407-7417.2002

75. Minson AC, Hodgman TC, Digard P, Hancock DC, Bell SE, Buckmaster EA. An analysis of the biological properties of monoclonal antibodies against glycoprotein D of herpes simplex virus and identification of amino acid substitutions that confer resistance to neutralization. J Gen Virol. 1986;67: 1001–1013. doi:10.1099/0022-1317-67-6-1001

76. Gierasch WW, Zimmerman DL, Ward SL, VanHeyningen TK, Romine JD, Leib DA. Construction and characterization of bacterial artificial chromosomes containing HSV-1 strains 17 and KOS. J Virol Methods. 2006;135: 197–206. doi:10.1016/j.jviromet.2006.03.014

77. Blanchet CE, Spilotros A, Schwemmer F, Graewert MA, Kikhney A, Jeffries CM, et al. Versatile sample environments and automation for biological solution X-ray scattering experiments at the P12 beamline (PETRA III, DESY). J Appl Crystallogr. 2015;48: 431–443. doi:10.1107/S160057671500254X

78. Graewert MA, Franke D, Jeffries CM, Blanchet CE, Ruskule D, Kuhle K, et al. Automated pipeline for purification, biophysical and x-ray analysis of biomacromolecular solutions. Sci Rep. 2015;5: 10734. doi:10.1038/srep10734

79. Wilkins MR, Gasteiger E, Bairoch A, Sanchez JC, Williams KL, Appel RD, et al. Protein identification and analysis tools in the ExPASy server. Methods Mol Biol Clifton NJ. 1999;112: 531–552. doi:10.1385/1-59259-584-7:531

80. Panjkovich A, Svergun DI. CHROMIXS: automatic and interactive analysis of chromatography- coupled small-angle X-ray scattering data. Bioinformatics. 2018;34: 1944–1946. doi:10.1093/bioinformatics/btx846

81. Franke D, Jeffries CM, Svergun DI. Correlation Map, a goodness-of-fit test for one-dimensional X-ray scattering spectra. Nat Methods. 2015;12: 419–422. doi:10.1038/nmeth.3358

82. Manalastas-Cantos K, Konarev PV, Hajizadeh NR, Kikhney AG, Petoukhov MV, Molodenskiy DS, et al. ATSAS 3.0: expanded functionality and new tools for small-angle scattering data analysis. J Appl Crystallogr. 2021;54: 343–355. doi:10.1107/S1600576720013412

83. Svergun DI. Determination of the regularization parameter in indirect-transform methods using perceptual criteria. J Appl Crystallogr. 1992;25: 495–503. doi:10.1107/S0021889892001663

84. Hajizadeh NR, Franke D, Jeffries CM, Svergun DI. Consensus Bayesian assessment of protein molecular mass from solution X-ray scattering data. Sci Rep. 2018;8: 7204. doi:10.1038/s41598-018-25355-2

85. Petoukhov MV, Franke D, Shkumatov AV, Tria G, Kikhney AG, Gajda M, et al. New developments in the ATSAS program package for small-angle scattering data analysis. J Appl Crystallogr. 2012;45: 342–350. doi:10.1107/S0021889812007662

86. Yang J, Yan R, Roy A, Xu D, Poisson J, Zhang Y. The I-TASSER Suite: protein structure and function prediction. Nat Methods. 2015;12: 7–8. doi:10.1038/nmeth.3213

87. Svergun D, Barberato C, Koch MHJ. *CRYSOL* – a Program to Evaluate X-ray Solution Scattering of Biological Macromolecules from Atomic Coordinates. J Appl Crystallogr. 1995;28: 768–773. doi:10.1107/S0021889895007047

88. Rueden CT, Schindelin J, Hiner MC, DeZonia BE, Walter AE, Arena ET, et al. ImageJ2: ImageJ for the next generation of scientific image data. BMC Bioinformatics. 2017;18: 529. doi:10.1186/s12859-017-1934-z

89. Schindelin J, Arganda-Carreras I, Frise E, Kaynig V, Longair M, Pietzsch T, et al. Fiji: an open- source platform for biological-image analysis. Nat Methods. 2012;9: 676–682. doi:10.1038/nmeth.2019

90. Valentini E, Kikhney AG, Previtali G, Jeffries CM, Svergun DI. SASBDB, a repository for biological small-angle scattering data. Nucleic Acids Res. 2015;43: D357–363. doi:10.1093/nar/gku1047

91. Receveur-Brechot V, Durand D. How random are intrinsically disordered proteins? A small angle scattering perspective. Curr Protein Pept Sci. 2012;13: 55–75. doi:10.2174/138920312799277901

