## Supplementary Material for "Herpes simplex virus 1 protein pUL21 stimulates cellular ceramide transport by activating CERT"

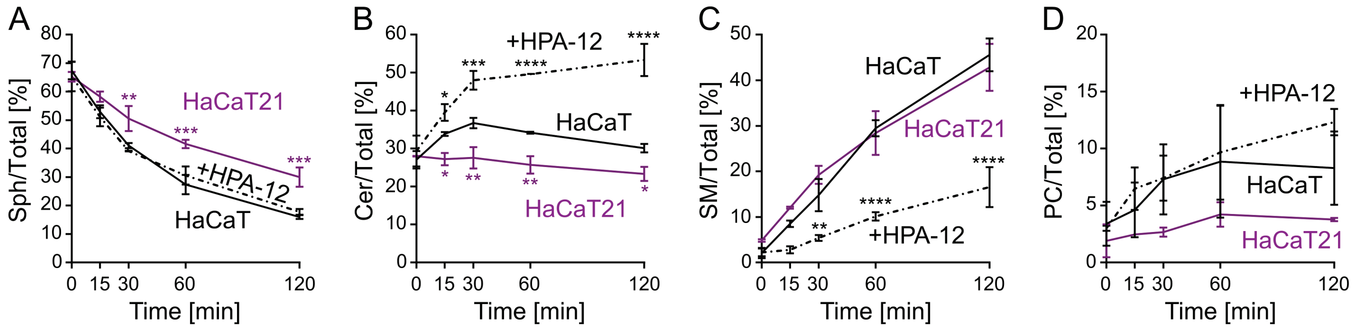


**Figure S1. HSV-1 pUL21 alters the rate of sphingosine metabolism in cultured cells.** Quantitation of the alkyne- (**A**) sphingosine (Sph), (**B**) ceramide (Cer), (**C**) sphingomyelin (SM) and (**D**) phosphatidylcholine (PC) intensities as percentage fraction of total signal. The data represents two independent experiments (mean ± SEM). Data points are labelled if significantly different to parental HaCaT cells: *, $p < 0.05$; **, $p < 0.01$; ***, $p < 0.001$; ****, $p < 0.0001$ (two-way ANOVA with Dunnett's multiple comparisons test).

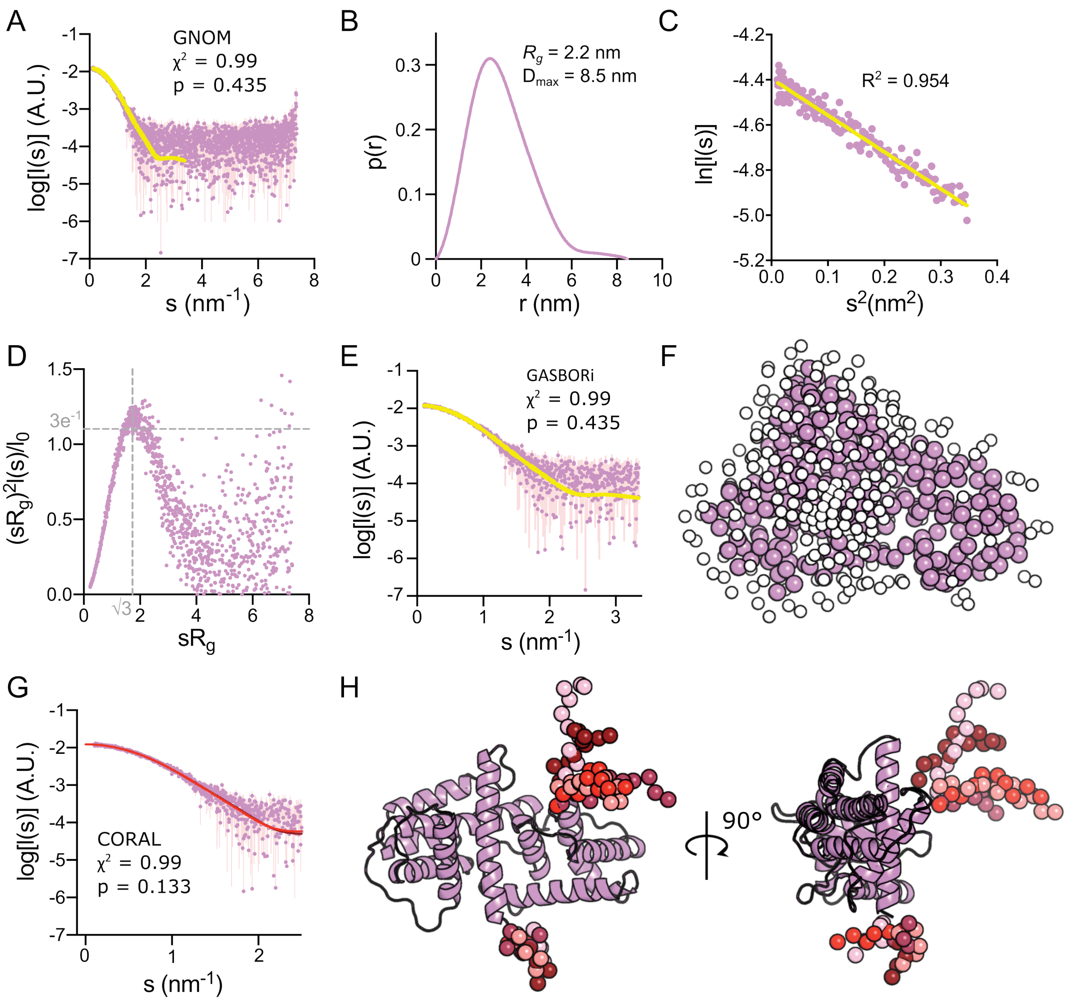


**Figure S2. pUL21C is a globular protein.** (**A**) SAXS profile for $H_6$-pUL21C. The reciprocal-space fit of the *p(r)* profile to the SAXS data is shown as a yellow line. $\chi^2$, fit quality; p, Correlation Map (CorMap) probability of systematic deviations between the model fit and the scattering data (Franke, Jeffries, and Svergun 2015). (**B**) The real-space distance distribution function, *p*(*r*), calculated from the SAXS profile. (**C**) The Guinier plot ($sR_g < 1.3$) is linear, representing an aggregate- and repulsion-free system. (**D**) Dimensionless Kratky plot. Grey dotted lines indicate the expected maximum of the plot for a compact protein ($sR_g = \sqrt{3}$, $(sR_g)^2I(s)/I(0) = 3e^{-1}$). (**E**) Fit of an *ab initio* dummy-residue model calculated using GASBOR to the SAXS profile. (**F**) GASBOR dummy-residue model. (**G**) Fit to SAXS profile of the five best pseudo-atomic models of $H_6$-pUL21C obtained by CORAL (continuous lines coloured in different shades of red, only topmost line is distinct because the lines overlay almost perfectly). $\chi^2$ and CorMap p value are shown for one representative fit. (**H**) CORAL pseudo-atomic model of $H_6$-pUL21C. pUL21C appears as violet ribbons and the regions modelled by CORAL are shown as spheres with colours corresponding to the fits shown in (**G**).

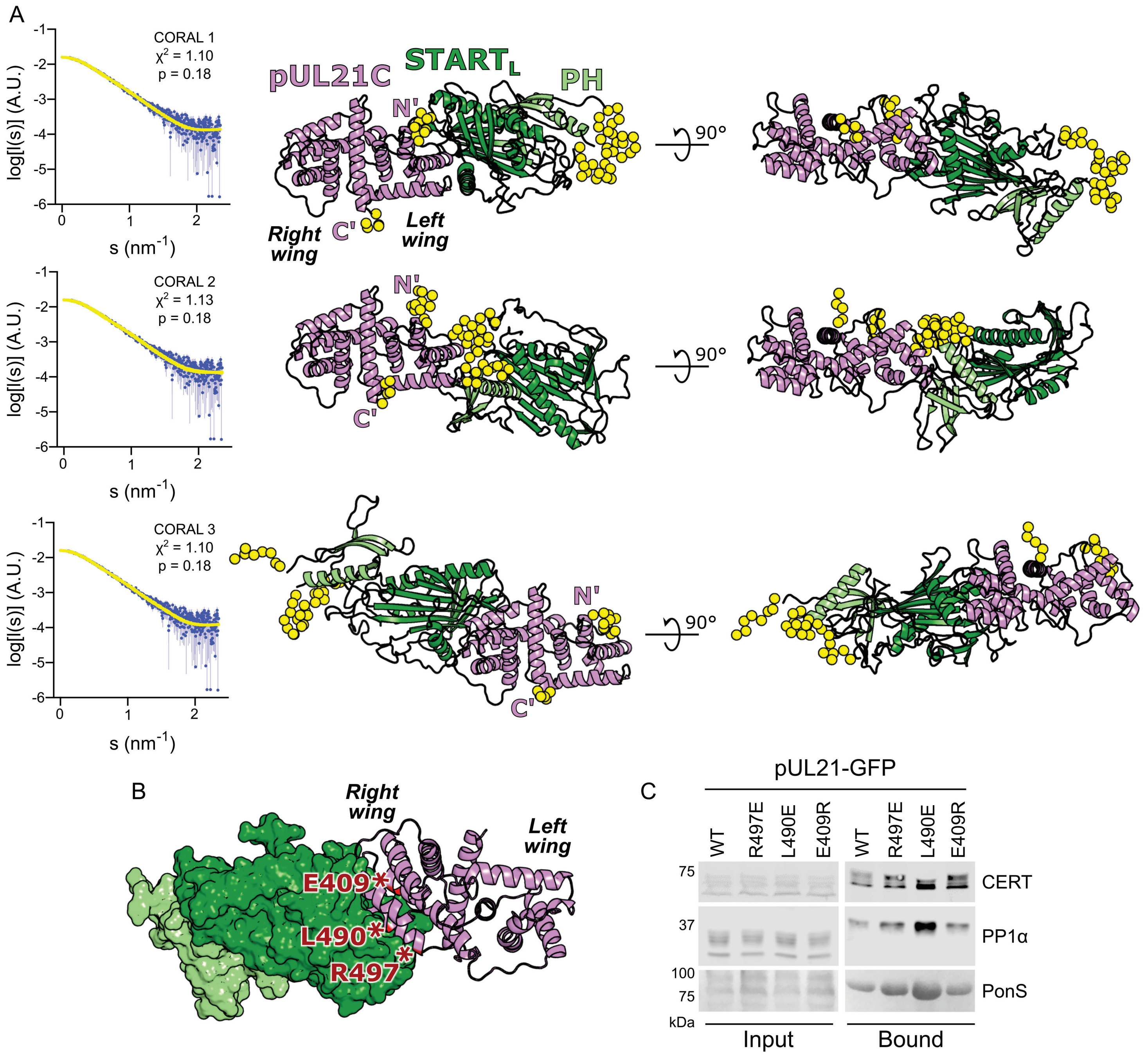


**Figure S3. Pseudo-atomic models of the H6-miniCERTL:H6-pUL21C complex.** (**A**) The best three pseudo-atomic models of the H6-miniCERTL:H6-pUL21C complex were selected based on their fit to the SAXS data (Fig 3D, lowest $\chi^2$). The fit of the computed scattering (yellow) to the H6-miniCERTL:H6-pUL21C SAXS profile for each model is shown, as are $\chi^2$ and CorMap p values. The pseudo-atomic models of the H6-pUL21C (violet cartoon) and H6-miniCERTL (PH, light green cartoon; STARTL, dark green cartoon) complex are shown in two orthogonal orientations. The left and right 'wings' of the dragonfly-like pUL21C fold (Metrick and Heldwein 2016) are labelled and the termini of the domain are marked. Regions absent from the crystal structures that were modelled by CORAL are shown as yellow spheres. *CORAL 1* (top) and *CORAL 2* (middle) represent the models shown in Figure 3, where the left wing of pUL21 binds miniCERTL, whereas in *CORAL 3* (bottom) the right wing of pUL21 binds miniCERT$_L$. (**B**) *CORAL 3* model of the $H_6$-miniCERT$_L$:$H_6$-pUL21C complex, with PH and START$_L$ domains of miniCERT$_L$ shown as light and dark green surfaces, respectively. pUL21 residues located at the hypothetical binding interfaces that were selected for further investigation are highlighted with red asterisks and labelled. (**C**) HEK293T cells were transiently transfected with GFP, wild-type (WT) pUL21-GFP or pUL21-GFP where residues at the putative miniCERT$_L$ binding surface of the *CORAL 3* model (**B**) were mutated. At 24 hours post-transfection cells were infected with ΔpUL21 HSV-1 (MOI = 5) and at 16 hours post-infection cells were lysed, tagged proteins were captured using GFP affinity resin, and the bound proteins were subjected to SDS-PAGE and immunoblotting using the antibodies listed. Ponceau S (PonS) staining of the nitrocellulose membrane before blocking is shown to confirm efficient capture of GFP-tagged proteins.

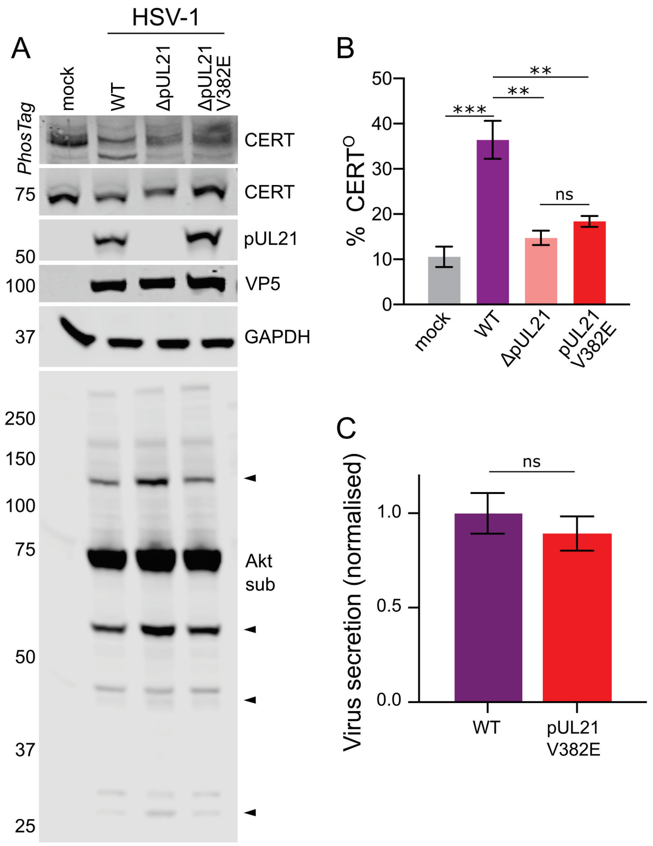


**Figure S4. pUL21-mediated dephosphorylation of CERT in Vero cells does not impact HSV-1 secretion.** (**A**) Vero cells were infected at MOI = 5 with wild-type (WT) HSV-1, HSV-1 lacking pUL21 expression (ΔpUL21), or a pUL21 point mutant virus ($pUL21^{V382E}$). Lysates were harvested at 16 hpi in the presence of phosphatase inhibitors and subjected to SDS-PAGE plus immunoblotting using the antibodies listed. Where indicated, the gel was supplemented with PhosTag reagent to enhance separation of CERT phosphoforms. The antibody recognising phosphorylated Akt substrates (Akt sub) illustrates activity of the HSV-1 kinase pUS3, several substrates of which are dephosphorylated in a pUL21-dependent manner (arrowheads) (Benedyk et al. 2021). VP5, infection control; GAPDH, loading control. (**B**) Quantitation of the CERT dephosphorylation level (ratio of $CERT^{O}$ to total CERT) in cells infected with WT or mutant HSV-1, as determined by densitometry. Results are presented as mean ± SEM from three independent experiments. One-way ANOVA with Dunnett's multiple comparisons test was used for the statistical analysis (ns, non-significant; **, $p < 0.01$; ***, $p < 0.001$). (**C**) Virus release into the culture supernatant from Vero cells infected with WT or mutant HSV-1 at MOI = 10. Samples were harvested at 12 hpi and virus infectivity in the cells versus the culture medium was measured by titration on Vero cells. The fold change in secretion of infectivity into the culture medium for $pUL21^{V382E}$ versus WT HSV-1 is shown as mean ± SEM of one independent experiment performed in technical triplicate.

**Table S1. SAXS parameters.**

| | $H_6$-miniCERT$_L$ | $H_6$-pUL21C | $H_6$-miniCERT$_L$: $H_6$-pUL21C |
|---|---|---|---|
| ***Data-collection parameters*** | | | |
| Radiation Source | Petra III (DESY, Hamburg, Germany) | | |
| Beamline | EMBL P12 | | |
| Detector | DECTRIS Pilatus 6M | | |
| X-ray wavelength (nm) | 0.124 | | |
| Sample-to-detector distance (m) | 3.0 | | |
| Temperature (°C) | 20 | | |
| Exposure time (s), Data frames (#) | 0.1, 47 | | |
| Measured protein concentrations (mg/mL) | 1.73–6.91 | 0.68 | NA |
| Injected protein concentration (mg/mL) | NA | NA | 4.3 |
| Measured *s*-range ($nm^{-1}$) | 0.07–7.219 | 0.08–7.37 | 0.05–7.36 |
| Final working *s*-range ($nm^{-1}$) | 0.13–4.52 | 0.10–3.35 | 0.08–5.08 |
| ***Structural parameters*** | | | |
| $I(0)$ (a.u.*) [from $p(r)$] | $0.024 \pm 1.5\times10^{-5}$ | $0.012 \pm 6.4\times10^{-5}$ | $0.016 \pm 5.9\times10^{-5}$ |
| Real-space $R_g$ (nm) [from $p(r)$] | 2.7 | 2.2 | 3.5 |
| $I(0)$ (a.u.*) (from Guinier) | $0.024 \pm 1.7\times10^{-5}$ | $0.012 \pm 5.9\times10^{-5}$ | $0.016 \pm 4.6\times10^{-5}$ |
| $R_g$ (nm) (from Guinier) | 2.66 | 2.21 | 3.31 |
| $D_{max}$ (nm) | 9.05 | 8.5 | 13.6 |
| Porod volume estimate (Vp, $nm^3$) | 70.5 | 11.7 | 85 |
| *Molecular-mass ($M_r$) determination* | | | |
| $M_r$ from Bayesian consensus (kDa) | 46 | 32 | 56 |
| $M_r$ credibility interval (kDa) | 43–47 | 30–33 | 47–58 |
| Expected $M_r$ from sequence (kDa) | 44.9 | 29.3 | 73 |
| ***Software employed*** | | | |
| Primary data reduction | *SASFLOW* | | |
| Data processing | *PrimusQT/GNOM(5.0)* | | |
| *Ab initio* analysis | *GASBOR(2.3.i)* | | |
| Computation of model intensities | *CRYSOL(2.8.3)* | | |
| Pseudoatomic modelling | *CORAL (1.1)* | | |
| ***Small Angle Scattering Biological Data Bank*** | | | |
| SASBDB accession codes | *SASDNC7* | *SASDNB7* | *SASDND7* |

*a.u. Arbitrary units
